# Anticipatory organization of neural population dynamics speeds behavioral decisions

**DOI:** 10.64898/2026.06.30.735699

**Authors:** Julia C Gorman, Tim Sainburg, Trevor S McPherson, Timothy Q Gentner

## Abstract

Expectations guide behavior and shape sensory responses in single neurons, but their influence on population-level neural dynamics is unknown. Here, we employ a dynamical systems framework to examine the collective spiking activity of neuronal populations in the auditory forebrain of European starlings, a species of songbird, as they categorize natural song syllables while sensory expectations are manipulated. We show first that sensory-driven neural population spiking activity traces smooth, low-dimensional latent trajectories that closely reflect the identity of sensory signals. Like the stimulus-driven responses of single neurons, the geometry of the population trajectories is also modulated by expectation. In single neurons, expectation sharpens differences between responses to signals in the same category, but at the population-level the effect is opposite: expectation increases the similarity between responses to signals in the same category. To understand how population-level response dynamics can differ from those in single neurons, we develop (and test empirically) a dynamical model that relates spiking activity at these two biological scales. The model leverages response redundancy between neurons, a capacity we term degeneracy-enabled remapping, and enables the observed simultaneous expectation-dependent increases in the separability of single-neuron responses *and* decreases in the separability of population trajectories in the task-potent subspace, i.e., the population activity dimensions tied to behavioral categorization. Examining the relationship between expectation-modulated population trajectories and behavior in detail, we find that single-trial categorization errors are tied to drift in the trajectory toward the opposing task-potent manifold. This suggests that expectations help establish structured, hypothesis-dependent initial conditions that precede the target-driven population response. In support of this, both behavioral accuracy and behavioral reaction time are predicted by the direction of early population motion within the task-potent subspace. We conclude that expectation drives anticipatory organization of population response variability into a structured, behaviorally relevant geometry that pre-positions subsequent population activity on task-potent manifolds to support rapid, accurate, behavioral outcomes.

## 1 INTRODUCTION

Perception changes continuously over time, reflecting ongoing neural processes that integrate incoming sensory evidence with internal states and behavioral goals [1, 2]. A central challenge is to understand the neurobiological structure of these ongoing processes. Classical approaches emphasize firing rates and receptive fields in single neurons, treating perception and sensory signals as discrete events and relying on trial-averaged responses [1]. These methods reveal how some individual neurons respond to stimuli, but are not well-suited to characterizing collective population activity over time. Population-level analyses instead track how collective activity organizes into low-dimensional time-varying trajectories in latent neural state space, reflecting ongoing internal processes that single-neuron measures may obscure [2–5]. In motor systems, these trajectories demonstrate how intentions and actions emerge from coordinated activity [6–8]. Applying the same tools to sensory systems has proven harder, however, in part because perception often lacks the clear behavioral outputs that anchor motor analyses. As a result, the link between sensory population activity and perception is not well-understood [4].

Population-level activity is not simply an average over constituent neurons, but can provide access to qualitatively different levels of response description [3, 5]. For example, the variance around trial-averaged single-neuron responses may be shared across multiple neurons, and this shared variance may itself possess a stable structure across trials in which any single neuron participates only intermittently [9, 10]. At the single-neuron level, this structured variability appears as noise. Critically, this population covariance is not uniformly distributed across neural state space. Rather, it can occupy specific subspaces whose alignment (or misalignment) with task-relevant directions determines whether it is consequential for behavior [11–13]. These findings reinforce the idea that perceptual states are reflected in not just in the mean trajectory of neural activity, but in how that trajectory is constrained from trial to trial [1]. One idea is that the fluctuations across trials reflect meaningful internal processes shaped by task demands, uncertainty, attention, and prior experience, which correlate with behavioral outcomes including accuracy and behavioral response latency [14– 16].

Here we consider one such potential constraint, expectation, shapes sensory population trajectories moment-to-moment and across trials. We ask whether sensory population dynamics in songbird pallial auditory populations are organized by expectations, whether this organization requires active task engagement, and whether it predicts relevant behavioral outcomes. Songbirds offer a powerful model system for these questions. Their auditory systems are highly specialized for processing acoustically complex vocal signals, and the auditory caudomedial nidopallium (NCM) and caudomedial mesopallium (CMM) integrate information over extended timescales, represent learned auditory categories, and receive both bottom-up sensory input and top-down feedback [17–22]. Field L is a more primary auditory region providing major ascending input to these areas [20–22]. Using a perceptual classification task in which natural vocal signals cue categorical expectations about upcoming vocalizations [23], we combine large-scale neural recordings in awake, behaving animals across the avian auditory forebrain with dimensionality reduction and dynamical systems modeling to track and understand population trajectories over time. We show that sensory expectation organizes the geometry of population trajectories in ways that reflect the animal’s categorical behavior across trials, and behavioral accuracy and reaction time on single trials.

## 2 RESULTS

### Expectation biases auditory categorization behavior

We trained European starlings (n = 10) using an operant procedure to categorize conspecific song syllables drawn from nine separate synthesized syllable continua (Fig. 1A; materials and methods). Each continuum spanned the naturalistic acoustic space, in 128 smoothly varying steps, between two distinct, natural, starling song syllables (Fig. 1B) [23, 24]. We set the midpoint of each continuum as the arbitrary categorical boundary between the two endpoint syllables (referred to as category ‘A’ and ‘B’) and trained subjects to associate syllables on each side of the boundary with different operant responses. The subjects’ task on each trial was to assign a randomly selected syllable from one of the nine continua (the ‘target’) to either category A or B by pecking either the left or right response port following stimulus presentation (Fig. 1A). Targets near the continuum endpoints were easy to classify and those near the category boundary were harder to classify. Once subjects reached a 70% accuracy on target syllable categorization alone, we introduced predictive syllables (‘cues’) that preceded the target on a subset of trials [25]. Cue syllables provided probabilistic information about the likely category (either A or B) of the upcoming target on a given trial. We used a range of cues during training and behavioral testing to maintain stimulus control by the target syllable (materials and methods), but for subsequent analyses we focus on the two cues that accurately predicted the correct target category (i.e., were valid) on ∼80% of trials in which they occurred (Fig. 1C), and inaccurately predicted the target category (i.e., were invalid) on ∼20% of trials in which they occurred (Fig. 1C).

**Figure 1:**
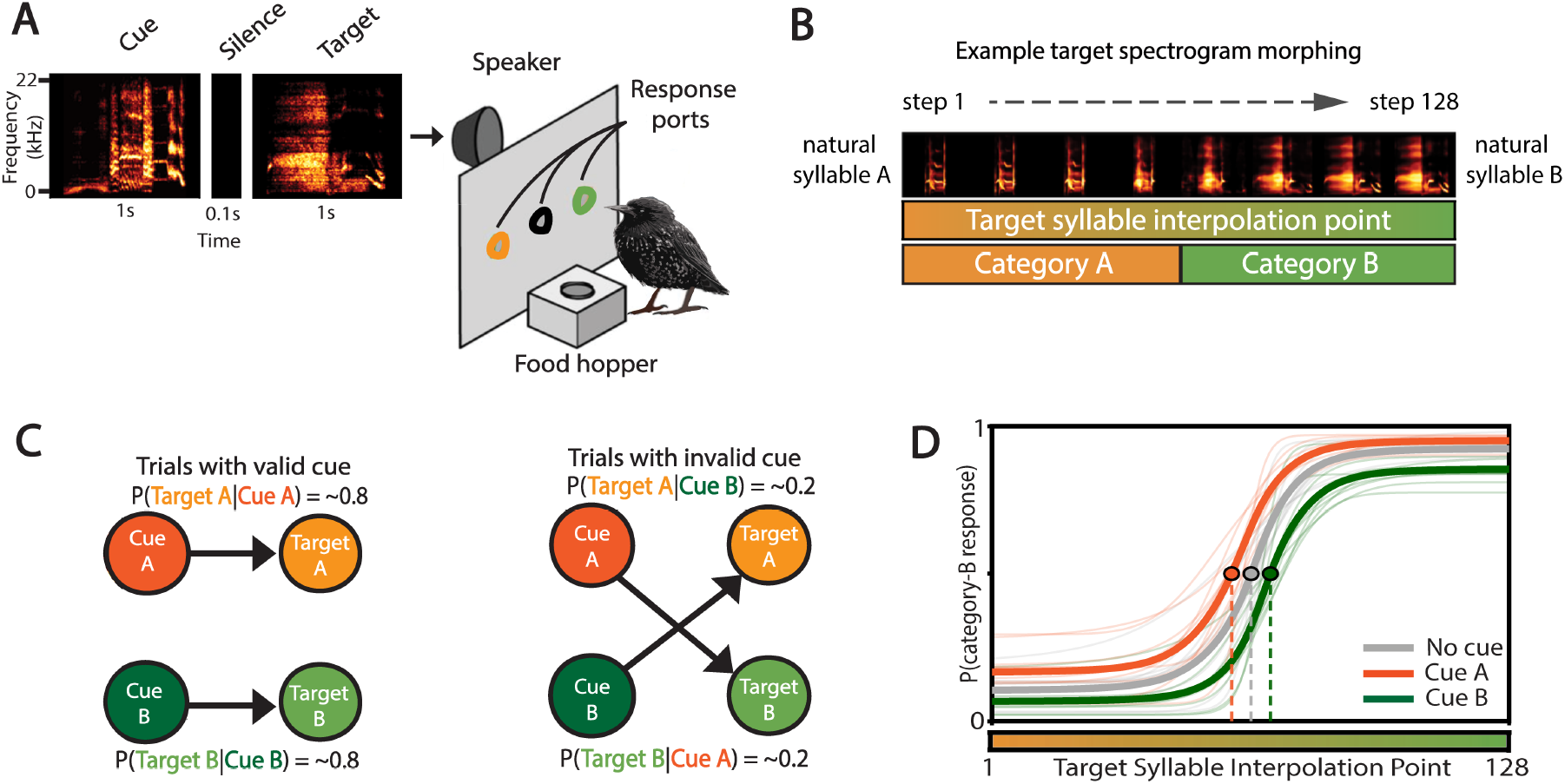
Behavioral paradigm. **(A)** Spectrograms showing an example stimulus sequence, and operant apparatus. On each trial subjects could hear two song syllables separated by a short silence (on left). The first syllable (the ‘cue’) indicated the likely category membership for the second stimulus (the ‘target’). Subjects indicated their choice for the categorical membership of the target at the end of the stimulus sequence by pecking the left or right response port (outlined in orange and green) to indicate either category A or B, respectively. The cue only appeared on a subset of trials. **(B)** Schematic of one target syllable continuum out of nine. Spectrograms (top) showing 8 1-s long sample syllables evenly spaced across the smoothly varying 128-syllable synthetic continuum (middle, schematized by graded coloration) that morphed natural syllable ‘A’ into natural syllable ‘B’. The mid-point of the continuum was arbitrarily set as the category A/B boundary (bottom). **(C)** Cued trial schematic. One of two possible cue syllables (cue-A or cue-B, orange or green) preceded the target syllable on 78% of analyzed trials. On *∼*80% of cued trials, the cue provided a valid prediction of the category membership for the following target (left). On *∼*20% of cued trials, the cue provided an invalid prediction of the category membership for the following target (right). **(D)** Behavioral performance. Psychometric curves showing the probability of a category-B response across an example target syllable continuum for the no cue, cue-A, and cue-B conditions. Thin lines show per subject curves, bold lines give the mean across subjects (bold; *n* = 10).

Under assumptions of Bayesian inference, the cue syllable functions as a categorical prior and so should shift the decision boundary toward the cued category; the more strongly the subject weights the cue, the larger the shift (relative to uncued trials) in the psychometric midpoint [26, 27]. The starlings trained to classify syllables along these continua show smooth, monotonic psychometric functions (Fig. 1D; per-continuum psychometric fits are shown in Fig. S1), with midpoints centered near the categorical boundary. As previously reported [23], the subjects’ behavior is well-modeled by Bayesian integration. Consistent with this, we find that valid cues shift the midpoint of the psychometric function toward the cued category relative to trials without a cue (Fig. 1D; cue-A: Δ*T*_50_ = 6.29 ± 1.13 steps, *W* = 0, *p* = 2.0 × 10^*−*3^; cue-B: Δ*T*_50_ = −4.70 ± 0.99 steps, *W* = 2, *p* = 5.9 × 10^*−*3^; *n* = 10 subjects, two-sided paired Wilcoxon) . Per-subject fit parameters, trial counts, and median reaction times are reported in Table S1. Thus, manipulating expectation, in this case using a predictive song syllable, biases categorization of future vocal elements.

### Expectation differentially modulates target-driven single-neuron and population-level responses

To understand how expectation acts at neurophysiological levels, we recorded extracellular spiking responses from multiple single neurons throughout the auditory pallium in a subset of subjects (*n* = 7) using 32- and 64-channel silicon probes (materials and methods), while subjects continued to perform the song-syllable-categorization task. We extracted a total of 7,524 single units across 225 different recording days, organized into populations of 10 to 170 simultaneously recorded neurons in Field L, the caudomedial mesopallium (CMM), and the caudomedial nidopallium (NCM; Fig. 2A; see Table 1). In the absence of histological confirmation of recording sites, and because probes frequently spanned the borders between these adjacent regions, units could not be assigned with certainty to a single area; we therefore pool across regions for all analyses and report per-region counts (Table 1) for sampling only. All neural analyses examine activity during the target-stimulus window exclusively as we are specifically interested in how the prior cue reshapes the neural response to the target, not the response to the cue itself (Fig. 2B). Previously we showed that for single neurons in these regions recorded under similar behavioral conditions, expectation increases the precision of the sensory driven response, such that responses to acoustically similar targets in the cue category become more separable [23]. To test whether this result holds in the present dataset, we smoothed single-neuron spike trains with a Gaussian kernel to produce firing-rate vectors (termed spike vectors; Fig. 2C) [23], then compared these spike vectors across cue conditions. To enable acoustically matched comparisons across cue conditions, we grouped the 128 interpolation steps within each of the nine continua into four bins, yielding 36 target syllable strata per population. We then computed pairwise cosine similarity (CS) within each stratum, separately for trials with valid or invalid cues (Fig. 2D).

**Table 1:** Dataset summary. Behavioral and electrophysiological dataset used in this study. *Behavior* summarizes all 10 trained subjects and their behavioral trials. The electrophysiology columns summarize the 7 subjects implanted for recording (*All regions*) and the breakdown by recording region (NCM, CMM, Field L). Because several subjects were recorded in more than one region, the per-region subject counts exceed the 7 unique implanted subjects: each subject is counted in every region from which it contributed populations. Neural populations and units reflect the inclusion criteria for the mixed-effects analyses (*≥* 10 isolated single units per neural population and *≥*2 neural populations per recording site, after exclusions). *Units (range)* and *Units (mean)* give the per-population single-unit yield, and *Prin. components* the effective dimensionality entering the population analyses. *Total trials* and its sub-rows count trials in the included neural populations; the cue (valid / invalid / no-cue), block (active / passive), and outcome (correct / incorrect) categories are not mutually exclusive (a single trial is, e.g., valid, active, and correct at once), so the sub-rows are not additive. Valid / invalid / no-cue partition trials by cue condition; active / passive by block type; correct / incorrect apply to responded (active) trials.

|  | Behavior | All regions | NCM | CMM | Field L |
| --- | --- | --- | --- | --- | --- |
| Subjects | 10 | 7 | 3 | 3 | 2 |
| Recording days | 2,750 | 225 | 144 | 47 | 34 |
| Probe locations | — | 13 | 8 | 3 | 2 |
| Units (range) | — | 10–170 | 10–170 | 10–102 | 10–111 |
| Units (mean) | — | 37.8 | 36.9 | 42.1 | 35.6 |
| Prin. components | — | 7 | 8 | 6 | 7 |
| Total trials | 2,434,924 | 887,434 | 605,069 | 189,101 | 93,264 |
| Valid | 1,533,278 | 165,831 | 104,092 | 35,070 | 26,669 |
| Invalid | 370,832 | 90,705 | 63,714 | 11,349 | 15,642 |
| No-cue | 530,814 | 630,898 | 437,263 | 142,682 | 50,953 |
| Active | — | 175,143 | 97,369 | 51,807 | 25,967 |
| Passive | — | 712,291 | 507,700 | 137,294 | 67,297 |
| Correct | — | 149,147 | 79,456 | 48,013 | 21,678 |
| Incorrect | — | 25,996 | 17,913 | 3,794 | 4,289 |

**Figure 2:**
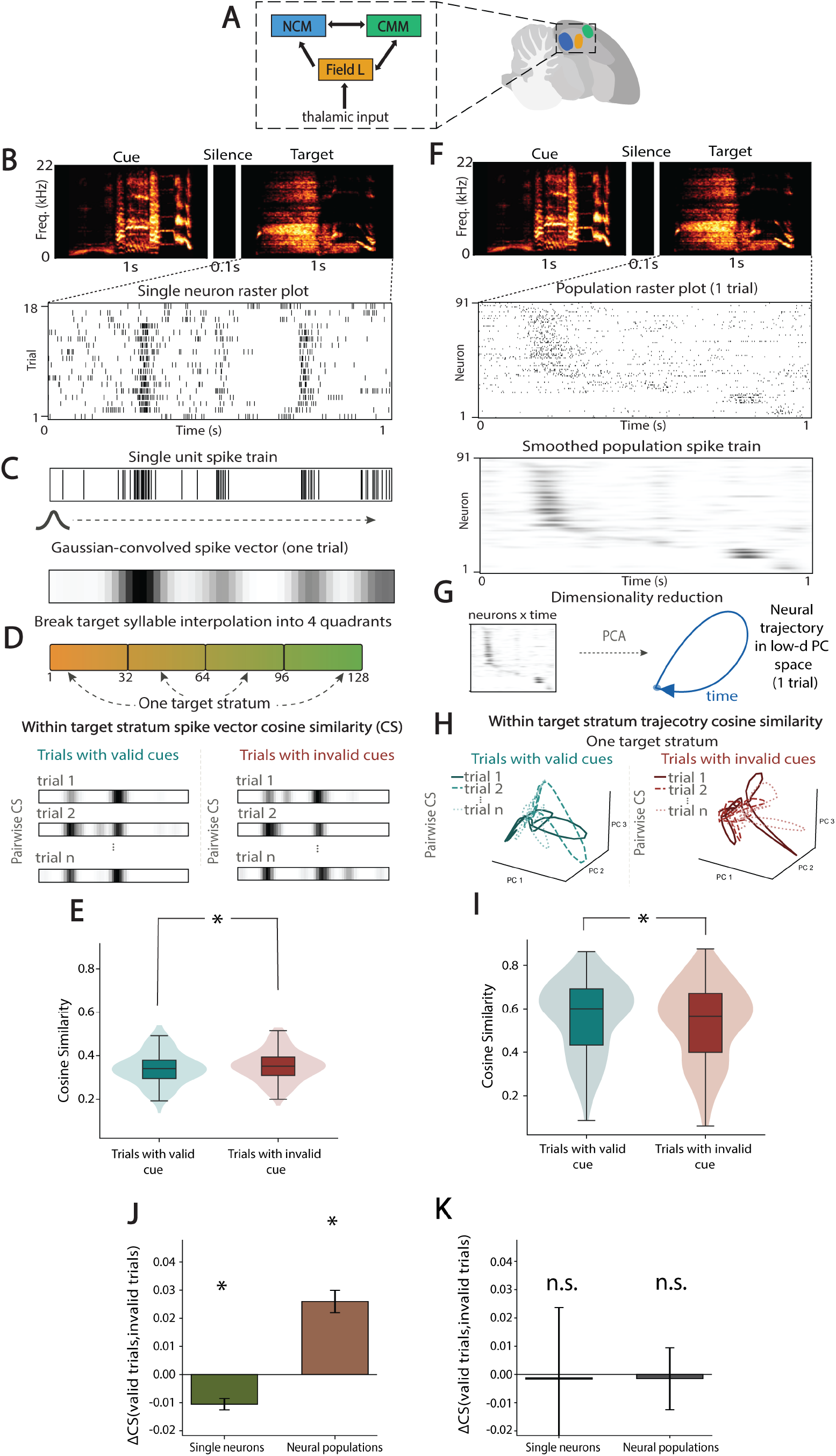
Cue validity has different effects on stimulus encoding at the single-neuron and population levels. **(A)** Recording regions. Electrode arrays targeted Field L (primary auditory region), the caudomedial mesopallium (CMM), and the caudomedial nidopallium (NCM). **(B)** Single-neuron recording schematic. Cue, silence, and target syllables are presented sequentially; spike times from a single unit are displayed as a raster across trials. All neural analyses examine the target-stimulus window only. **(C)** A single-unit spike train (top) is convolved with a Gaussian kernel to produce a smooth spike vector for one trial (bottom). **(D)** Within-target-stratum spike-vector CS schematic. Pairwise cosine similarity (CS) is computed between Gaussian-smoothed spike-rate vectors within acoustically matched trial groups (strata), shown separately for trials with a valid or invalid cue. **(E)** Distribution of pairwise spike-vector cosine similarity between single-neuron responses to acoustically matched (same-stratum) targets, shown separately for trials with a valid and invalid cue (*n* = 201 neural populations from 7 subjects). Each violin shows the full distribution across neural populations; the overlaid box marks the interquartile range, the central line the median, and the whiskers the remainder of the distribution. The bracket and * denote a significant difference between valid and invalid cues. **(F)** Population recording schematic. Spectrograms (top) are followed by a population raster for a single trial (*N* neurons time; top) compiled into a smoothed population spike-train as an *N×T* matrix (bottom). **(G)** Dimensionality reduction. The *N×T* smoothed population spike-train matrix is projected via PCA into a low-dimensional space, yielding a neural trajectory through principal-component space for each trial (time flows along the trajectory). **(H)** Within-target-stratum trajectory CS schematic. Pairwise CS is computed between population trajectories within each stratum for trials with a valid or invalid cue. **(I)** Within-category trajectory cosine similarity of population responses, shown separately for trials with a valid and invalid cue (*n* = 201 neural populations from 7 subjects). Violins, box, median, and whiskers as in (E); the bracket and * denote a significant difference between valid and invalid cues. **(J)** Mean ΔCS (valid invalid) for single neurons and neural populations (*n* = 201 neural populations from 7 subjects). Bars show mean *±*SEM; * denotes significance from 0. **(K)** Mean ΔCS (valid *−* invalid) during passive listening; the difference is not significantly different from zero (single neurons *n* = 108 neural populations from 5 subjects; populations *n* = 109 neural populations from 6 subjects). Error bar shows mean *±* SEM.

Consistent with prior observations [23], we find that expectation modulates single neuron responses. Specifically, pairs of responses to target syllables within the same stratum that are preceded by a valid cue have significantly lower cosine similarity than those preceded by an invalid cue (Fig. 2E; *n* = 201 neural populations from 7 subjects; linear mixed-effects model [LME] with a by-subject random intercept, valid − invalid *β* = −0.0090, 95% CI [−0.0160, −0.0021], *p* = 1.1 × 10^*−*2^). Thus, expectation sharpens the response of individual neurons to acoustically matched (same-stratum) syllables. As noted previously, this is consistent with heightened perceptual acuity, but does not help to explain categorization behavior directly, which should intuitively be tied to an *increase* in the response similarity to syllables within the same stratum/category.

A neural response aligned with categorization behavior may be observed in the downstream targets of these forebrain single neurons, or in the non-independent collective activity of these same neurons. To test the latter possibility, we constructed smoothed population spike vectors for simultaneously recorded units (materials and methods; Fig. 2F) and applied Principal Component Analysis (PCA) to project population activity into a low-dimensional latent space, yielding trial-aligned neural trajectories (Fig. 2G) [28]. We first asked whether these trajectories reflected the identity of the target syllable. We find that a simple cross-validated linear decoder (materials and methods) using the single-trial population trajectory is able to recover which of the nine continua, and which stratum on that continua, the target belongs to, at rates significantly better than label-shuffled data (continuum identity: *β* = +0.120, 95% CI [+0.078, +0.162], *p* = 2.3 × 10^*−*8^, *n* = 214 neural populations; continuum bin: *β* = +0.105, 95% CI [+0.075, +0.135], *p* = 6 × 10^*−*12^, *n* = 224 neural populations; Δ balanced accuracy above chance, LME with subject as random intercept, 7 subjects). Thus, like the single neurons response, the population activity trajectories carry stimulus specific information.

To understand whether expectation modulates these population trajectories, we computed the pairwise CS between trajectories within each stratum, separately for trials with valid or invalid cue conditions (Fig. 2H). Again we find that expectation modulates the response, however, the effect of cue validity at the population level is the opposite of that in single neurons. Valid cues increase the CS between within-category trajectories, while invalid cues decrease it (Fig. 2I; *n* = 201 neural populations from 7 subjects; *β* = +0.024, 95% CI [+0.012, +0.036], *p* = 1.1 × 10^*−*4^). That is, the same conditions that sharpen response differences in single neurons yield more similar responses at the population level. The scale-dependent difference between single-neurons and populations is directly summarized in the ΔCS comparison (Fig. 2J) showing that, relative to invalid cues, valid cues lead to a decrease in CS between target syllable responses at the single-neuron level and an increase in CS at the population level. Thus, unlike the single neuron responses, the population trajectories align with the animals’ behavior suggesting that categorical representations are encoded in the collective activity of the auditory forebrain.

The foregoing results are consistent with the conclusion that population-level responses carry expectation-dependent, behaviorally relevant structure that is not visible in single-neuron responses where expectation sharpens tunings. But expectation implies a top-down modulation that cannot be entirely explained by bottom-up stimulus driven effects. Although our analyses only consider the single-neuron and population responses to matched target syllables, cue syllables preceding targets on trials with a valid and invalid cue differ, leaving open the possibility that stimulus differences account for the observed effects. To examine this possibility, we manipulated task engagement by presenting cue-target syllable sequences (identical to those used in the categorization behavior) during passive epochs in which the house-light was extinguished and the apparatus was inoperable. If the foregoing effects are due to expectation rather than pure stimulus drive, they should abate or disappear when the task demands are removed. Indeed, we find that the cue-dependent organization of population geometry requires active task engagement. When subjects hear the same cue-target sequences during passive playback, the increase in within-category trajectory similarity under valid cues is abolished, and target responses produce indistinguishable population trajectories and single-neuron responses regardless of cue validity with a valid−invalid difference that did not differ from zero across neural populations (Fig. 2K: single neurons *β* = −0.0006, 95% CI [−0.0248, +0.0237], *p* = 0.96, *n* = 108 populations from 5 subjects; populations *β* = −0.0015, 95% CI [−0.0125, +0.0095], *p* = 0.79, *n* = 109 populations from 6 subjects) The full per-condition distributions of within-category cosine similarity during passive playback are shown for both scales in Supplementary Fig. S2. This collapse in the cue-dependent population-level effect without active engagement rules out an exclusively bottom-up, stimulus-driven response.

The scale-dependent effects of expectation motivate the remainder of the study: we next develop a latent-dynamics model to ask what circuit mechanism could produce simultaneous single-neuron sharpening and population-level consolidation, and then examine whether the shifts in population geometry have a functional payoff for behavior.

### A latent-dynamics model of degeneracy-enabled remapping reproduces the scale-dependent difference

To understand the observed differences in how expectation shapes single-neuron and population-trajectory responses to categorized song syllables, we built a minimal generative latent-dynamics model (Materials and Methods) that captures the key components of our data and yields quantitative predictions. In the model, a low-dimensional latent state *x*_*t*_ summarizes the population’s internal state and evolves in time according to:

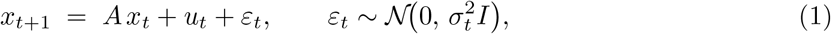

where *A* is a fixed linear dynamics matrix, *u*_*t*_ is the external drive from the target syllable at time *t*, and *ε*_*t*_ is Gaussian noise with time- and condition-dependent variance 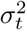 (Materials and Methods). Population responses are generated by a time-varying linear readout *r*_*t*_,

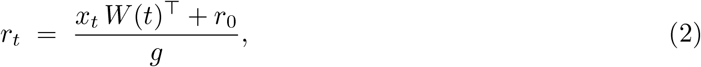

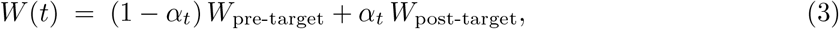

where *r*_0_ is a positive baseline that ensures non-negative firing rates and *g* is a scalar gain that uniformly rescales activity (both held fixed across trials and conditions; neither affects the geometric analyses, which operate on mean-centered activity). *W* (*t*) is a readout matrix that maps the low-dimensional latent state into the higher-dimensional neural activity space; its rows specify how each neuron weights the latent variables and are not constrained to be mutually orthogonal. The mixing coefficient evolves as

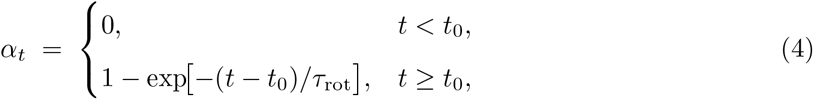

so that prior to target onset at *t*_0_ the readout equals *W*_pre-target_ and, after onset, rotates smoothly toward a post-target readout *W*_post-target_ with time constant *τ*_rot_. We refer to this within-trial rotation of *W* (*t*) from *W*_pre-target_ to *W*_post-target_ as *remapping*. The endpoint of this rotation, *W*_post-target_, is itself drawn per trial from a cue- and target-conditioned distribution (Materials and Methods), so the same within-trial rotation produces different reorganizations of single-neuron firing patterns across cue conditions and trials.

We then let *u*_task_ be the unit-norm direction in activity space along which category-A and category-B mean responses differ, and decompose the readout into components aligned with and orthogonal to *u*_task_,

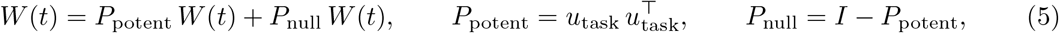

where *W*_null_ ≡ *P*_null_ *W* (*t*) and *W*_potent_ ≡ *P*_potent_ *W* (*t*) for the two components. Rotations confined to *W*_null_ change neuron-wise activity patterns while leaving the projection of population activity onto *u*_task_ essentially unchanged. Whether remapping alters the *categorization-relevant component* of the population response depends on whether the weight changes are concentrated in *W*_null_, which reorganizes activity in directions orthogonal to *u*_task_ and therefore preserves the projection that supports categorization, or in *W*_potent_, which alters that projection and so changes the information available for categorizing targets as either category A or category B. The model structure reflects a basic property of population codes; namely, that neurons share variability. This produces low-rank covariation and enables dimensionality reduction, of course [2, 5, 29], but also means that many distinct patterns of single neuron activity can give rise to the same low-dimensional latent population trajectory [30, 31]. Our model controls this redundancy by changing the neuron-wise weights, which we refer to as *degeneracy-enabled remapping*. It enables cues to set distinct pre-target population state conditions, *x*_0_, that reflect category-A or category-B expectations. Importantly, these initial conditions are not arbitrary: a category-A cue places *x*_0_ closer to the category-A side of the task-potent subspace, so that the pre-target population state is aligned with the correct categorization geometry when the target arrives. The weight changes that accomplish this readjustment reside almost entirely in *W*_null_, leaving the task-potent trajectory geometry intact. This model yields three testable predictions, which we evaluate first through simulation, validating the scale-dependent empirical differences shown in Fig. 2, and then test directly in our recordings.

#### Prediction 1: Cues set distinct pre-target initial states

The model predicts that cue-conditioned initial states should bias latent trajectories before the target arrives. Because cues initialize different latent states, pre-target trajectory segments should cluster by cued category and then diverge along validity-dependent paths once the target arrives. During the 100ms window prior to target onset, cues already separate the initial, pre-target segments of the trajectories in the low-dimensional activity subspace, placing the population closer to either the category-A or category-B manifold before any target-driven activity occurs (Fig. 3A; *n* = 100 trials per cue condition). Once the target arrives, simulated example trajectories for a category-A and category-B target follow distinct paths through PCA space (Materials and Methods): valid-cue trials take a shorter path toward the correct categorization manifold, whereas invalid-cue trials follow longer, more dispersed trajectories (Fig. 3B; *n* = 10 trials per stimulus condition) [8, 32–34].

**Figure 3:**
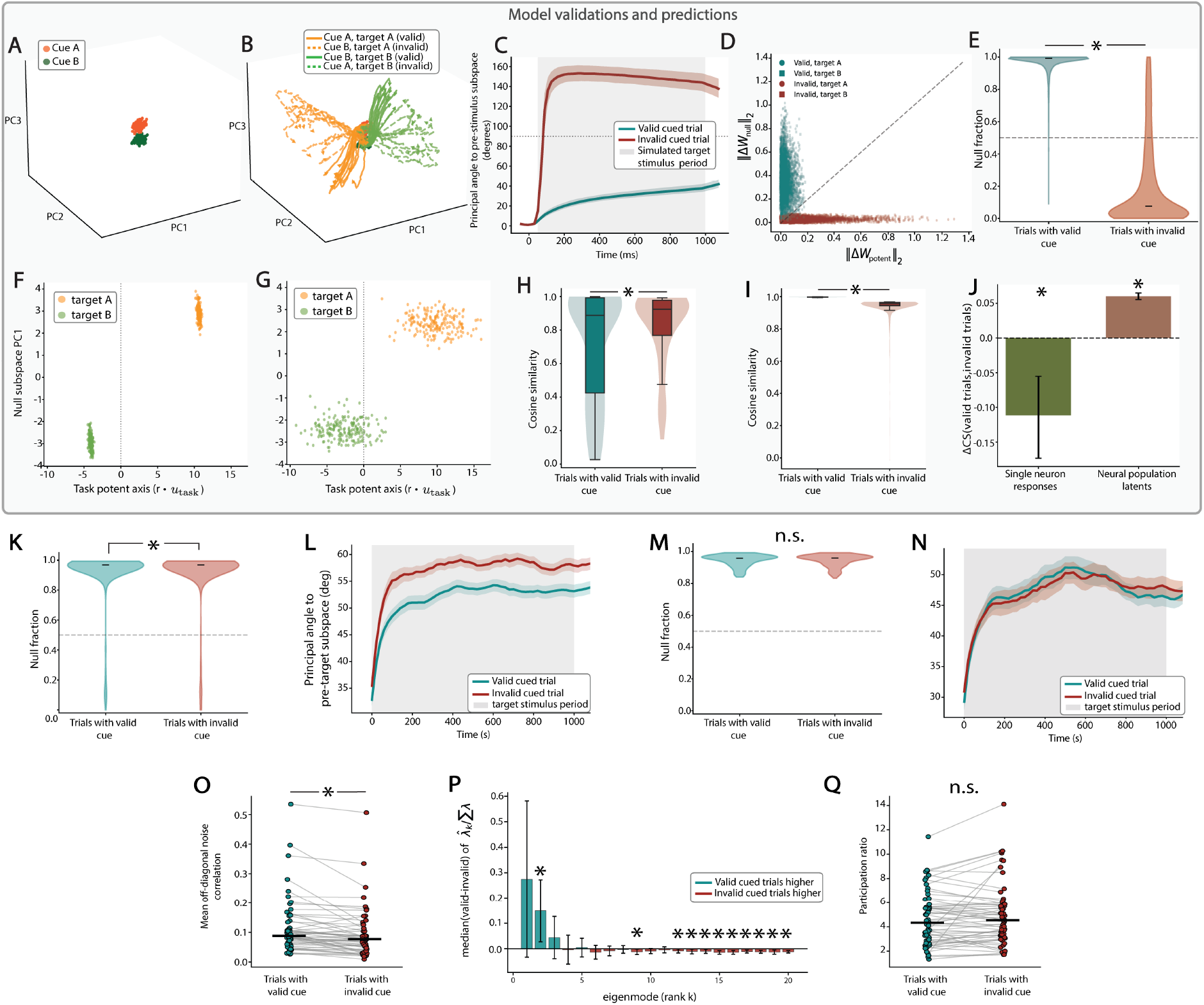
A latent-dynamics model of degeneracy-enabled remapping: simulated predictions and empirical validation in the auditory forebrain. Panels A–J show model simulations; K–Q, auditory-forebrain recordings. **(A)** Three-dimensional PCA of simulated pre-target trajectory segments from the 100 ms before target onset, colored by cue identity (100 trials per cue). **(B)** 3D PCA of simulated population trajectories for category-A and category-B targets across the full trial epoch. Trajectories are colored by target category, with solid lines for trials with a valid cue and dashed lines for an invalid cue (*n* = 10 trials per stimulus condition). **(C)** Per-trial trajectory rotation, the angle between the pre-target anchor and the current state *x*_*t*_ in mean-centered activity, for trials with a valid or invalid cue (*n* = 2400; 1200 each). Lines show medians, shading the IQR, and grey the target epoch. **(D)** Per-neuron weight displacement (*W*_pre-target_ *→ W*_post-target_) split into task-potent (*x*) and task-null (*y*) components. Each dot is one neuron on one trial (*n* = 10,000); color indicates cue validity, shape target identity, and the dashed line equal potent and null displacement. **(E)** Per-neuron null fraction for the same *n* = 10,000, split by cue validity. Each violin shows the full distribution and the black bar the median; the dashed line marks 0.5, and the bracket and * a significant valid*−*invalid difference (medians: valid *≈* 1.000, invalid *≈* 0.031). **(F)** Single-trial stimulus-window activity on trials with a valid cue, projected onto the task-potent axis (*x*; *r u*_task_) versus the leading null dimension (*y*) and colored by target category (*n* = 200 trials per target). **(G)** As (F) for trials with an invalid cue. **(H)** Pairwise spike-vector cosine similarity of simulated single-neuron responses for trials with a valid or invalid cue (*n* = 100 neurons). Each violin shows the distribution, the box the IQR, and the line the median; the bracket and * denote a significant difference. **(I)** Pairwise trajectory cosine similarity of simulated population trajectories for trials with a valid or invalid cue (*n* = 1200 trials per condition); glyphs as in (H). **(J)** Mean valid *−* invalid difference in trajectory CS (population) and spike-vector CS (single-neuron). Bars show the mean *±*SEM and * significance from zero (population, *n* = 1200 trials per condition; single-neuron, *n* = 100 neurons). **(K)** Null-subspace fraction of stimulus-window displacement on active trials, for trials with a valid or invalid cue (*n* = 58 populations, 7 subjects). Violins show the distributions, bars the medians, and the dashed line 0.5. **(L)** Mean principal angle between the pre-target and target-epoch subspaces (top *k* = 3 PCs) for trials with a valid or invalid cue; shading shows *±*SEM (*n* = 58 populations; 53 paired). **(M)** As (K) for passive playback (*n* = 18 populations, 4 subjects). **(N)** As (L) for passive playback (*n* = 22 populations, 5 subjects). **(O)** Mean off-diagonal noise correlation per population for trials with a valid or invalid cue (paired; black line, mean; *n* = 55 populations, 7 subjects). **(P)** Cross-validated eigenspectrum (cvPCA): per-mode LME estimate of the valid *−* invalid difference in variance-normalized eigenvalues (*n* = 58 populations, 7 subjects). Error bars show 95%CIs and markers FDR-corrected significance. **(Q)** cvPCA participation ratio for trials with a valid or invalid cue (paired; black line, mean; *n* = 58 populations, 7 subjects). Each line is one population, colored by whether the ratio was higher on trials with a valid or an invalid cue.

#### Prediction 2: Cue validity sets the magnitude of remapping

The model predicts that the magnitude of rotation required to reach the correct categorization geometry depends on cue validity. An invalid cue initializes *x*_0_ near the opposing categorization manifold, generating a large angular displacement between *W*_pre-target_ and *W*_post-target_, and thus requiring a substantial *W*_potent_ component to realign to the correct categorization geometry. For valid cues, *x*_0_ initializes near the correct manifold, with a smaller angular displacement and a reduced *W*_potent_ component (Fig. 3C; stimulus-window rotation: valid median = 32.7^*°*^, IQR [29.0^*°*^, 36.7^*°*^]; invalid median = 136.9^*°*^, IQR [125.8^*°*^, 146.3^*°*^]; Δ median = +104.2^*°*^, 95% CI [+103.0^*°*^, +105.4^*°*^]; Mann–Whitney *U* = 1.44 × 10^6^, *p <* 10^*−*100^, *r* = 1.0; *n* = 2400 simulated trials, 1200 valid and 1200 invalid). Thus the larger *W*_potent_ component produced by an invalid cue requires more substantial alteration of the target-driven trajectory.

#### Prediction 3: Valid-cue remapping concentrates in the null subspace

Finally, the model predicts that the single-neuron weight changes which implement the population-level remapping are concentrated in the null subspace when the cue is valid. This is the key prediction of our framework; namely, that reorganization of single-neuron responses confined to the null subspace leaves the behaviorally relevant, population-level target-driven response unchanged. To test this directly, we decompose the single neuron weight displacements into their null and potent components,

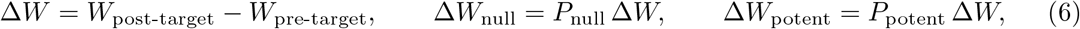

and ask how the magnitudes of each component compare neuron-by-neuron and trial-by-trial. If valid cues remap activity by organizing null-subspace population variance, then for a single neuron on trials with a valid cue the null component should be greater than the potent component and the opposite should be true on trials with an invalid cue. The per-neuron scatterplot of null-vs-potent components confirms this: for single neurons, components on 100.0% of trials with a valid cue fall above the unity line, indicating null-dominant displacement, compared with only 15.5% of neurons on invalid trials, whose displacements are predominantly directed along the potent axis (Fig. 3D; *n* = 10,000 per-neuron observations from 100 simulated trials). We quantify this separation with a per-neuron “null fraction” index (Materials and Methods), which equals 1 when a neuron’s displacement lies entirely in the null subspace and 0 if it lies entirely in the potent subspace. We show that neurons in our model have null fractions close to 1 on trials with a valid cue, and close to 0 on trials with an invalid cue (Fig. 3E; valid median = 1.000, IQR [1.000, 1.000]; invalid median = 0.031, IQR [0.007, 0.184]; Δ median = +0.969, 95% CI [+0.966, +0.971]; Mann–Whitney *U* = 2.46 × 10^7^, *p <* 10^*−*100^, rank-biserial *r* = +0.999; *n* = 10,000 per-neuron observations). The same null/potent split is evident at the level of single-trial population activity: projecting each trial’s mean stimulus-window activity onto the task-potent axis and the leading null-space dimension, valid-cue trials form compact, well-separated category clusters along the task-potent axis with their trial-to-trial variability confined to the null subspace (Fig. 3F), whereas invalid-cue trials instead spread along the task-potent axis itself, degrading the separation between categories (Fig. 3G; *n* = 200 trials per target, of 600 simulated per condition).

#### Simulations reproduce both scale-dependent effects

When null-biased remapping is the organizing principle, the latent population trajectory can remain aligned in the task-potent subspace even as changes unfold in constituent single neurons, as long as those changes lie in an orthogonal null subspace (Supplementary Fig. S3). We show widely different single-neuron response patterns project onto the same task-potent population readout on valid trials, whereas on invalid trials the same readout is dispersed. This reconciles our observation of expectation-induced decreases in the similarity of single-neuron responses and increased similarity of the simultaneous population dynamics. Indeed, the simulations reproduce both effects within a single population. At the single-neuron level, responses to within-category targets become more distinct following valid than invalid cues (Fig. 3H; Δ_V*−*I_ = −0.111, SEM = 0.030; *t*(99) = −3.50, *p* = 4.3 × 10^*−*4^; Wilcoxon *W* = 3029.0, *p* = 4.2×10^*−*2^; *n* = 100 simulated neurons). While at the population level, trajectories on trials with valid cues are more similar than those on trials with invalid cues, measured in the population 3D PCA subspace (Fig. 3I; Δ_V*−*I_ = +0.062, SEM = 0.003; Mann–Whitney *U* = 1.44 × 10^6^, *p* = 1.1 × 10^*−*94^, *r* = +1.0; *n* = 1200 simulated trials per cue condition). The mean valid−invalid difference at each scale is summarized side by side in Fig. 3J. The model thus provides a mechanistic account linking single-neuron heterogeneity to stable, low-dimensional trajectories that support target categorization and reproduces the scale-dependent differences observed empirically.

#### Null-biased remapping in the auditory forebrain

Having established that our model simulations reproduce the scale-dependent difference in the effects of expectation, we next test geometric predictions of the null-biased remapping model directly in our electrophysiological data. Specifically, we ask (i) whether valid cues concentrate single-neuron weight displacements in the population-level null subspace, and (ii) whether cue validity is tied to the magnitude of the subspace rotation and effective dimensionality of population-level dynamics.

#### Null-biased reorganization of target-evoked population activity

The model predicts that valid cues reorganize trial-to-trial activity within the null subspace while leaving the task-potent readout intact. The per-neuron weight remapping that produces this in the model is not observable in spiking data, so we test its direct consequence at the level of population activity. We define a task-potent direction in the smoothed population spike-train activity space. Within each neural population, we fit a linear decoder to discriminate category-A versus category-B target syllables using target-epoch mean activity, pooling trials across valid and invalid cues. For each trial we take the target-window population activity relative to its pre-target baseline, a time-by-neuron matrix, and split this population displacement into the component along the decoder axis (potent) and the orthogonal remainder (null). We summarize each trial by its null fraction, the share of the displacement’s total squared magnitude lying in the null subspace, computed over the whole population rather than per neuron. This linear-decoder axis serves as an empirical analog to the model’s task-potent axis *u*_task_, allowing us to test the null-subspace prediction directly in neural data. We observe that both conditions show predominantly null-subspace displacement which is expected given that the null space vastly outnumbers the single potent dimension [7]. However, the null fraction is significantly larger on trials with a valid cue than on trials with an invalid cue (Fig. 3K; *n* = 58 neural populations from 7 subjects; *β* = +0.0047, 95% CI [+0.0001, +0.0094], one-sided *p* = 0.022). As observed for population-level trajectory effect (Fig. 2K), this null-biased reorganization was contingent on active task engagement. Applying the same decomposition to passive playback trials, the cue-dependent effect on null-subspace displacement is absent, and trials following a valid or invalid cue show comparable null fractions (Fig. 3M; *n* = 18 neural populations from 4 subjects; *β* = +0.0001, 95% CI [−0.0009, +0.0011], *p* = 0.90). Thus, null-biased remapping can restructure single-neuron responses while preserving population structure in the task-potent subspace, but only when the animal is actively engaged in the categorization task.

#### Valid cues reduce population subspace rotation

One potential benefit of null-biased remapping is to pre-position the population in anticipation of the upcoming target. We tested this by asking how much the dominant dimensions of population activity reorient from the pre-target period to the target epoch, the population-geometry analog of the readout rotation in our model. For each neural population and cue condition we defined a pre-target subspace S_pre-target_ and a target-epoch subspace S_target_ as the top *k* = 3 principal components of the trial-averaged population activity within each window, and measured their reorientation as the mean canonical principal angle between them. Consistent with the model prediction, trials with an invalid cue rotate more from the pre-target period to the target epoch than trials with a valid cue (Fig. 3L; *β*_invalid*−*valid_ = +5.67^*°*^, 95% CI [+4.11^*°*^, +7.23^*°*^], *p* = 1.1 × 10^*−*12^, *n* = 58 neural populations from 7 subjects). As with the null-subspace reorganization, this rotation difference required active task engagement: during passive playback the pre-target-to-target rotation did not differ between valid and invalid cues (Fig. 3N; *β* = −0.15^*°*^, 95% CI [−1.26^*°*^, +0.95^*°*^], *p* = 0.79; paired Wilcoxon *p* = 0.58, median Δ = −0.03^*°*^; *n* = 22 neural populations from 5 subjects). This effect is preserved across a range of PCA components (*k* = 2–6; Supplementary Fig. S4). This supports the notion that expectation drives anticipatory organization of the population subspace that then constrains future stimulus driven activity.

#### Null-biased remapping reshapes population covariance

Beyond rotation, null-biased remapping makes a specific prediction about the covariance structure of population activity. If trial-to-trial remapping concentrates in shared null directions, neurons undergoing similar null-space fluctuations should co-vary more on trials with a valid cue than those with an invalid cue, because the null-space remapping weights under valid cues are larger [35, 36]. Consistent with this, we observe that mean pairwise noise correlations are significantly higher during valid-cued targets than during invalid-cued target syllables (Fig. 3O; *n* = 55 neural populations from 7 subjects; *β* = +0.0225, 95% CI [+0.013, +0.032], *p* = 5.2×10^*−*6^). Moreover, because a more correlated noise covariance is lower-rank by construction, concentrating variance into fewer leading eigenmodes, the increased noise covariance on validly cued trials should leave a clear spectral fingerprint. The cross-validated eigenspectrum (cvPCA [4]) confirms this prediction. Relative to invalid-cue trials, valid-cue trials carry more variance in the leading eigenmodes (Fig. 3P; mode 2: *β* = +0.015, FDR-corrected *p* = 4.8 × 10^*−*2^; mode 1: *β* = +0.027) and less in higher-order modes (Fig. 3P; modes 15–20: *β <* 0, FDR-corrected *p <* 0.05). Notably, the participation ratio does not differ significantly between conditions (Fig. 3Q; *n* = 58 neural populations from 7 subjects; *β* = −0.30, 95% CI [−0.69, +0.08], *p* = 0.12), indicating that valid cues reorganize *which* dimensions carry variability rather than compressing overall dimensionality. This is what null-biased remapping predicts: structured co-fluctuation in the null space, invisible to the task-potent readout.

### Expectation-driven pre-target geometry predicts single trial decision speed

Having established null-biased remapping as a useful organizing principle to understand the geometry of stimulus-driven sensory population responses, we next ask how this reorganization relates to behavioral outcomes on individual trials [37, 38]. Our latent-dynamics model treats population dynamics as explicitly tied to task goals, with expectation acting as a top-down organizing signal that constrains how stimulus-driven collective population activity moves through the task-potent subspace.

If expectation organizes population geometry along a task-potent decision axis, cues that support faster, more accurate decisions should advance the population further along this axis toward the manifold tied to correct categorization. To test this prediction within each neural population, we defined a task-potent decision axis by fitting a linear decoder that discriminated category-A from category-B target conditions using target-epoch activity (trained on no-cue trials and applied to cued trials; Materials and Methods). To quantify distance on the task-potent axis, we scored each trial by its decoder margin signed toward the true target category. Valid-cue trials carried larger target-aligned margins than invalid-cue trials, after balancing within stimulus strata to remove stimulus-composition bias (Fig. 4A; *β*_valid*−*invalid_ = +0.046, 95% CI [0.032, 0.061], *p* = 1.8 × 10^*−*10^; *n* = 195 neural populations across 7 subjects). Thus, valid cues drive population activity further into the task-potent direction toward the region associated with the correct response. This same axis also predicts behavioral response errors. Scoring each trial by the sign of its margin relative to its true target category, incorrect trials fell on the opposing-category side of the axis more often than correct trials, again after balancing within stimulus strata (Fig. 4B; *β*_incorrect*−*correct_ = +0.041, 95% CI [0.028, 0.055], *p* = 1.4 × 10^*−*9^; *n* = 195 neural populations across 7 subjects). This provides a geometric interpretation of behavioral errors: incorrect trials preferentially drift into the opposing task-potent direction, consistent with misalignment of latent dynamics with the task-potent subspace associated with the intended behavioral response.

**Figure 4:**
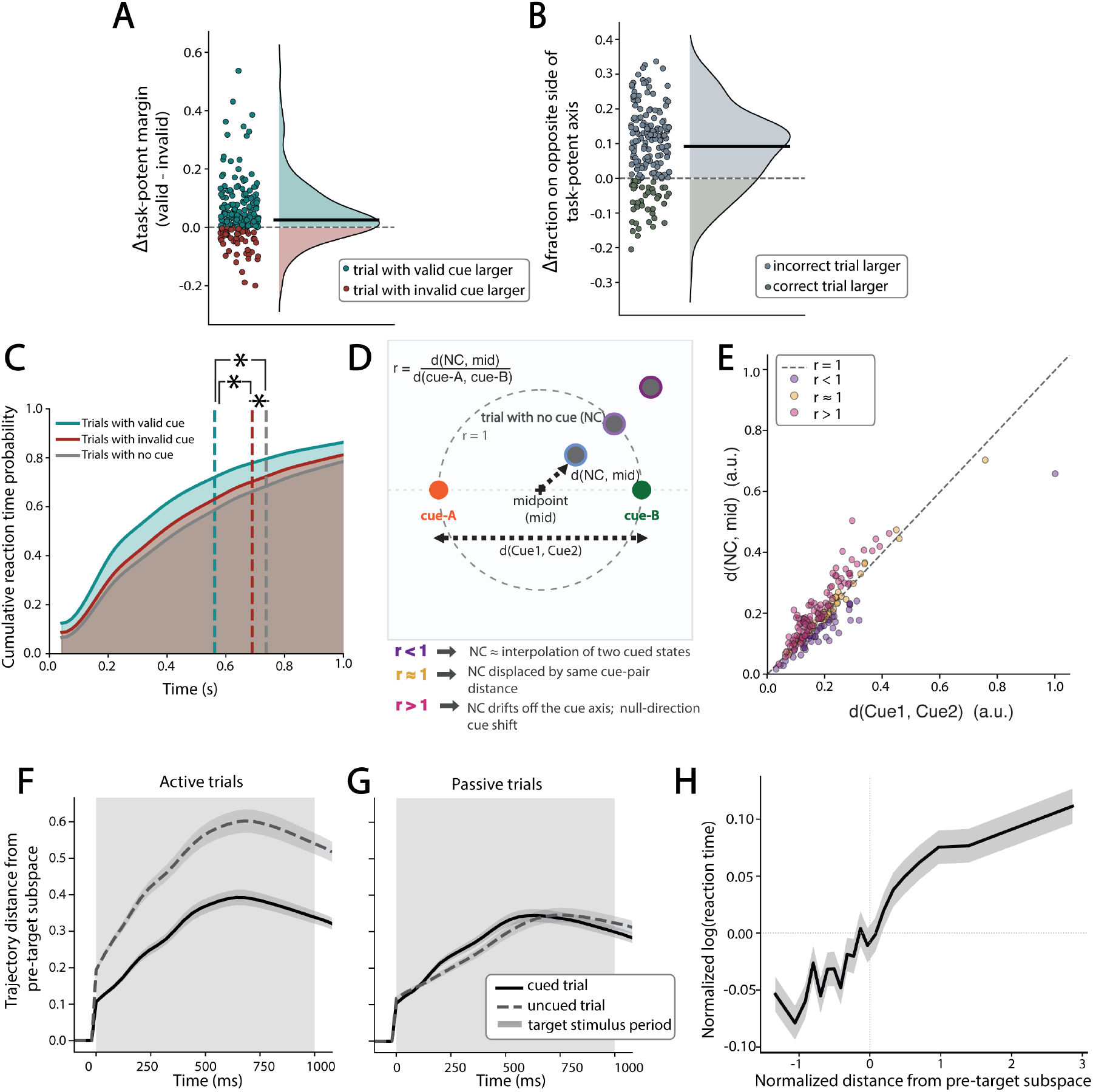
Cue-dependent population geometry requires task engagement and predicts the speed and accuracy of perceptual decisions. **(A)** Δ task-potent margin (valid *−* invalid) for each neural population. Each point is one population, colored by the sign of its difference (teal, valid *>* invalid; red, invalid *>* valid); the black bar marks the median (*n* = 195 populations, 7 subjects). **(B)** Δ wrong-side fraction (incorrect *−* correct) for each population on the held-out task-potent axis. Each point is one population, colored by the sign of its difference (grey, incorrect *>* correct; green, correct *>* incorrect); the black bar marks the median (*n* = 195, 7 subjects). **(C)** Cumulative distribution of reaction times for trials with a valid cue, an invalid cue, or no cue. Vertical dashed lines mark the median per condition (valid, 0.328 s; invalid, 0.426 s; no cue, 0.485 s; *n* = 455,659, 122,227, and 201,920 trials, respectively, across 10 subjects). All pairwise differences are significant. **(D)** Schematic of the pre-target centroid geometry comparison. Pre-target centroids are computed separately for the category-A cue, the category-B cue, and trials without a cue. The ratio *r* = *d*(No Cue, mid)*/d*(cue A, cue B) compares the displacement of the no-cue centroid from the cue-defined midpoint to the separation between the cue-conditioned centroids. *r <* 1 indicates the no-cue state lies between the two cued states; *r ≈* 1 indicates it is displaced from the midpoint by about the cue-pair distance; *r >* 1 indicates it drifts off the cue axis. **(E)** Scatter of *d*(No Cue, mid) versus *d*(cue A, cue B). Each point is one neural population, colored by the ratio *r* = *d*(No Cue, mid)*/d*(cue A, cue B) (purple, *r <* 1; gold, *r ≈* 1; magenta, *r >* 1). Points above the dashed unity line (*r* = 1) have the no-cue pre-target state farther from the cue-defined midpoint than the cue-conditioned centroids are from each other (*n* = 198 populations from 7 subjects).**(F)** Normalized Euclidean distance between the running target-epoch mean population vector and the pre-target mean, for cued (solid) and uncued (dashed) trials during active listening; distance is normalized by baseline pre-target dispersion. Shading shows *±* SEM (*n* = 198 neural populations from 7 subjects). **(G)** Same as (F) for passive listening (*n* = 109 neural populations from 6 subjects). The cue *×* modality interaction (Results) was tested on the 89 populations (5 subjects) recorded under both conditions; (F) and (G) show the full per-condition sets. **(H)** Distance from the pre-target subspace versus log reaction time, both z-scored within neural population and pooled across all active correct trials. The black line shows the mean log reaction time across 20 equal-frequency bins of the z-scored distance; shading shows *±* SEM (*n* = 89,232 trials).

Expectation also significantly affects decision speed, with per-subject median reaction times ordered valid *<* invalid *<* no-cue (Fig. 4C; Friedman *χ*^2^(2) = 14.25, *p* = 8.0 × 10^*−*4^, *n* = 10 subjects). A subject-aware trial-level mixed model confirmed the ordering. One way that cues could produce this behavioral speed-up is by pre-positioning the population at a more advantageous starting point in the task-potent subspace prior to target onset, so that less of the target-evoked population trajectory is spent moving toward the correct categorization manifold. This is what we mean by anticipation. To understand these ideas in more detail, we examine whether cues establish separable, hypothesis-aligned, pre-target initial conditions along the task-potent subspace, as predicted by our latent-dynamics model. We note that on trials without a cue, the population lacks an explicit categorical prior over target category, whereas on cued trials, both valid and invalid cues impose a categorical hypothesis before target onset. If expectation positions the population into a hypothesis-aligned initial condition, as we propose, then cued trials should establish opposing (category-A vs category-B), but structured, pre-target states, and the pre-target state on no-cue trials should sit off the hypothesis-defined axis set by the cued trials. Geometrically, we predict that the no-cue centroid should lie farther from the midpoint of the two cue-conditioned centroids than these two centroids are from each other (Fig. 4D). To test this, we define two cue-conditioned pre-target centroids from category-A and category-B cued trials for each population, and compare their separation to that of the no-cue centroid from the midpoint of the cue-conditioned centroids. Consistent with our predictions, the no-cue pre-target state lies farther from the cue-defined midpoint than these two states are from one another (Fig. 4E; *n* = 198 neural populations from 7 subjects; *β* = 7.9 × 10^*−*2^, 95% CI [1.4 × 10^*−*9^, 1.4 × 10^*−*8^], *p* = 1.0 × 10^*−*2^). This is consistent with the idea that expectation constrains the sensory population into a structured, hypothesis-dependent initial condition. To explain the slowest reaction times on uncued trials, we reasoned that in the absence of this anticipatory constraint the uncued population drifts off this hypothesis-driven axis entirely rather than settling between the two cued poles. To test this, we quantified pre-target drift as the normalized Euclidean distance between the running target-epoch mean population vector and the pre-target mean. Consistent with our prediction, correct no-cue trials drifted further from pre-target geometry than cued trials (Fig. 4F; *β* = −0.34, 95% CI [−0.39, −0.28]). During passive listening this difference in pre-trial initial conditions is abolished (Fig. 4G; cue × modality interaction *β* = +0.35, 95% CI [+0.28, +0.43], *p* = 4.3×10^*−*20^ ; *n* = 89 populations, 5 subjects with sufficient data). Finally, we ask whether the anticipatory organization of population geometry can be directly tied to faster reaction times on a trial-by-trial basis. Across nearly all of our neural populations, trials that start with a larger displacement from the pre-target subspace are associated with slower reaction times (Fig. 4H: LME *β* = +0.018, 95% CI [+0.015, +0.02], *p* = 2.0 × 10^*−*44^, *n* = 89, 232 trials, 132 neural populations from 7 subjects). This relationship holds across multiple measures of pre-target stability (pre-target variance, distance from population centroid; Supplementary Fig. S5), and is consistent with reaction time effects on cued trials alone: cued trials sit closer to pre-target geometry, and closer pre-target states predict faster responses (*β* = +0.0088, 95% CI [+0.006, +0.012], *p* = 5.8 × 10^*−*10^, *n* = 73, 104 trials, 132 neural populations from 7 subjects).

## 3 DISCUSSION

We demonstrate that sensory neural populations throughout the songbird auditory forebrain exhibit smoothly evolving, low-dimensional, trajectories that reflect coordinated spiking activity across the population. The trajectories are sensory-driven, reflecting the unique, immediate time-varying acoustics of different natural song syllables, but are reminiscent of the kinds of collective dynamics commonly observed in motor control regions in many species [6, 12, 39–41]. In addition, these trajectories are not fixed. Rather, expectations, unfolding over multi-syllable timescales, actively shape the geometry of sensory neuro-dynamics on a trial-by-trial basis. When animals are not actively categorizing song syllables, these expectation effects disappear, thus they are not solely produced by feed-forward stimulus drive. At the behavioral level, these rapid expectation-driven changes are tied to categorization accuracy and response speed on single trials.

We modulated expectation on each trial by presenting a pair of song syllables in which the first syllable (the cue) predicted the learned category to which the second syllable (the target) likely belonged. The cue orients the ongoing population activity so that it is biased toward the predicted, learned, categorical response at target onset. This orientation then constrains the initial trajectory of target-syllable-driven population activity to move along behaviorally potent dimensions. Acoustically different cues associated with different response categories initialize different population orientations. Thus, when a cue is valid, i.e., paired with a target syllable that falls into the predicted category, the target-driven trajectory evolves more quickly through the low-dimensional latent space and the animal responds faster. In contrast, an invalid cue, i.e. one paired with a target syllable in the opposing category, requires the initial target-driven trajectory to make a larger corrective rotation to move into the task-potent subspace. This leads to more variable trajectories and slower behavioral responses following invalid cues. We interpret this expectation-dependent initialization of the population geometry as an anticipatory neural process, whereby regularities in the environment structure ongoing population activity in anticipation of upcoming sensory stimuli and speed acquisition of organism-level behavioral goals [42–44]. In this sense, the system isn’t “representing” sensory stimuli. Rather, it has learned how to structure stimulus driven activity into a functional, behaviorally relevant subspace. This structured activity is a strong candidate for downstream modulation of processes underlying appetitive (pecking) and consummatory (feeding) behaviors.

Signatures of the observed anticipatory process reside at that population level. The dynamics and geometric organization of this activity are emergent properties of the underlying neural system and cannot be observed in single neurons. Indeed, we find that the same predictive cues that constrain population dynamics in the service of behavior also sharpen single-neuron responses, as shown previously [23]. That is, in the context of the current categorization task, expectation leads to *increased* within-category stimulus differentiability in single neuron responses and *decreased* within-category stimulus differentiability at the population level (Fig. 2J). These seemingly opposed effects at different biological scales seem paradoxical from the perspective where the population response is viewed as an average of the constituent single neurons. To understand this apparent paradox, we developed a latent-dynamics model that leverages the fact that the responses of single neurons within a population are not independent. As such, there exist many neuron-wise weight configurations that can theoretically yield the same low-dimensional population trajectory [30, 31]. We take advantage of this redundancy through a computational mechanism that we refer to as degeneracy-enabled remapping, whereby the network can reorganize single-neuron response covariance along dimensions that are irrelevant for, or null with respect to, task “readout”. This is similar to null-space models proposed for motor control [7, 11, 13, 45]. In our case, shifting stimulus-driven single-neuron covariance into behaviorally null dimensions enables dissociation of expectation-dependent sharpening in single-neuron selectivity and the convergence of population trajectories. We confirm predictions of our degeneracy-enabled remapping model in both simulation results and our empirical data (Fig. 3), and show that the stimulus-driven activity covariance is flexibly and functionally organized along behaviorally potent and null dimensions on a trial-by-trial basis.

The observed covariance signatures also distinguish our degeneracy-enabled remapping model from competing hypotheses tied to gain modulation. Uniform gain would increase separability on every scale at once and would not selectively elevate noise correlations or concentrate variance in the leading eigenmodes without changing overall dimensionality [35, 46, 47]. Instead, valid cues do both. They raise noise correlations and redistribute variance toward the leading modes while leaving the overall effective dimensionality (participation ratio) the same. This pattern of spectral organization is predicted by null-biased remapping rather than gain [1]. Overall, degeneracy-enabled remapping provides a powerful mechanism for neural systems to maintain distinct stimulus-driven responses in single neurons, while simultaneously organizing covariance between these same neurons to flexibly align with one or more behavioral goals based on top-down modulatory drive. The network level pattern of connectivity underlying the anticipatory reorganization of population activity can not be unequivocally determined from our data. We find that the scale-dependent effects are clearest in the secondary regions NCM and CMM, and weakest in the primary thalamo-recipient region Field L (Fig. S6). This is consistent with the idea that the reorganization draws on recurrent, feedback-rich circuitry [17, 19–22], but as with any extracellular data, the precise assignment of individual units within a population to any specific sub-field should be viewed cautiously. Moreover, relatively few subjects contributed data to any given region, and Field L in particular was represented by only two subjects, precluding a subject-level test. In addition, while our latent-dynamics model reproduces the scale-dependent geometry, it does so by imposing the null/potent organization by construction rather than deriving it directly from circuit connectivity, cell-type heterogeneity, or the source of the top-down cue signal. Future, explicit feedback- and cell-type-structured models should address this gap[48].

Although derivable from the same data, our measures of population activity differ from more classical measures of population responses, such as the average stimulus driven activity and the trial-by-trial neuron-wise response fluctuations, referred to as “stimulus” and “noise” correlations, respectively [1, 49]. Most importantly, both the stimulus and the noise correlation are typically pairwise metrics that summarize the distribution of spike count correlations across the population (and typically across time), and so average out structure in exactly which neurons co-vary with which other neurons at any point in time. Dimensionality reduction methods, such as the PCA used here, measure the time-varying patterns in this covariance directly across the population, by find-ing the axes along which activity co-varies, moment-by-moment and trial-by-trial. Thus, pairwise and population metrics reveal different attributes of the same neural activity [50]. For example, improved behavioral performance in cued visual attention tasks is often accompanied by a reduction in pair-wise noise correlations[49, 51]. In these detection and discrimination settings, global de-correlation improves performance because the suppressed shared variability lies along the same dimensions that carry stimulus information, where it would otherwise limit the recoverable signal [52]. Such an account may be sufficient when the behavioral goal is stimulus detection or discrimination, including in songbirds [19]. For tasks that require both generalization and discrimination, as is true for categorization, the capacity to confine correlated neural fluctuations to the behaviorally null subspaces without degrading the task-potent population activity may be highly efficient ways to maintain behavioral flexibility and use the same signals in the service of multiple behavioral goals.

Much of the debate over expectation in sensory systems is framed as a question about how and where representations of various components in a Bayesian decision process, such as the stimulus likelihood, prior, or posterior, are encoded [23, 53–58]. The present categorization behavior is well described by Bayesian integration across a range of cue reliabilities [23], and our physiological data can be interpreted within the general Bayesian framework, where what is encoded in the auditory forebrain depends on the scale at which one measures neural activity. At the single neuron level, responses sharpen, consistent with a change in the stimulus likelihood [23]. At the population level, spiking covariance is aligned to posterior-like categorization behavior in a way that analyses restricted to single-units cannot see [8, 11]. Focusing too closely on how these responses relate to (or encode) Bayesian probability distributions, however, misses an opportunity to understand what we contend is a more important aspect of the neural system’s behavior. Namely, that it has managed to align the expected target-driven activity subspace to the effective dimensions of the animal’s behavioral goals. Consistent with the sensory drive of action, our effects are tied to behavior in several important ways. As with a variety of other experience-dependent re-organization effects [59–64], the responses reported here require active engagement and disappear during passive listening (Figs. 2J,K, 4F,G). Likewise, expectation dependent re-organization is established prior to the target-driven neural trajectory (Fig. 3L), and on error trials the population drifts toward the opposing task-potent manifold (Fig. 4B) even for unambiguous target syllables, as if a misdirected anticipatory set steered activity toward the wrong goal. Like predictive coding and inference[56, 65], this anticipatory behavior acts to pre-configure the sensory population based on learned regularities in the environment[42, 43, 66]. Rather than an end product of altered sensory encoding, however, the neural response re-organization is channeled toward the behaviorally potent dimensions that drive action [67]. In this view the organization we describe is the sensory-side counterpart of the preparatory geometry of motor cortex [7, 45], where population activity is arranged along potent and null dimensions in advance of behavioral output. Our central finding is that stimulus-evoked activity within the sensory populations organizes along similar behavioral output dimensions.

## Acknowledgments

This work was supported by NIH grant 1F99NS141305 and a Center for Academic Research and Training in Anthropogeny (CARTA) fellowship to J.C.G., and by NIH grants R01DC018446 and R01DC018055, and NSF GRFP 2017216247 to T.Q.G. and T.S. We thank J. Xing, L. Stanwicks, and L. Ostrowski for their advice on the manuscript.

## Author Contributions

J.C.G. developed the geometric framework and analyses; T.S., T.S.M. and T.Q.G. designed experiments. T.S. and T.S.M. carried out experiments. T.Q.G. contributed to analysis conception and direction. J.C.G. and T.Q.G. wrote the paper; T.S. provided feedback on the manuscript.

## Data and Code Availability

All data and analysis code supporting the findings of this study will be made publicly available at https://github.com/juliagorman/anticipatory_geometry_paper upon publication.

## 4 MATERIALS AND METHODS

### Subjects

Subjects were adult European starlings (*Sturnus vulgaris*) both male and female, wild-caught and housed in a large mixed-sex aviary; 10 contributed behavioral data and 7 contributed neural data. During testing, subjects were held on a restricted-feeding schedule to optimize operant behavior and adjusted per subject to keep body weight within 10% of their ad-libitum baseline weight. Colony and operant chamber lighting tracked seasonal sunrise and sunset in San Diego, California. We did not otherwise control for photoperiod. All procedures were approved by the Institutional Animal Care and Use Committee of the University of California, San Diego under protocol S05383.

### Behavioral paradigm and training

We controlled subjects’ categorization behavior using an established 1-interval, two-alternative choice, operant task[23, 24]. The operant apparatus contained three response ports (left, center and right), a retractable food hopper, and a playback speaker all controlled by a custom Raspberry Pi-based system (PiOperant; https://github.com/gentnerlab/pyoperant.git). Target syllables were drawn from nine independent morph continua, each synthesized with a variational autoencoder trained on starling song acoustics that interpolated smoothly over 128 evenly spaced steps between two distinct, natural starling song syllables sampled from the full song repertoire [68] of a single conspecific male (Fig. 1D) [23, 24]. The two natural syllables at opposing endpoints of each continuum defined each category (labeled A or B) and were arbitrarily mapped to either the left or right response port, counterbalanced across subjects. Subjects initiated a trial with a peck at the center response port, which triggered the playback of a 1-sec long target syllable, after which the subject reported the category to which the syllable belonged by pecking either the left or right response port within a 5-sec window. Correct responses were reinforced with access to food on a variable-ratio schedule requiring a run of two to four correct responses (set per subject), and incorrect responses triggered a 5-sec timeout during which the house-light extinguished, food was inaccessible, and no new trial could be initiated. The inter-trial interval was a minimum of 1 second.

Training proceeded in stages. Subjects first learned to discriminate the two natural endpoint syllables of a single continuum. Once proficient, additional endpoint pairs were added until performance on all 18 endpoint syllables (9 continua × 2 endpoints) was significantly above chance. Subjects were then transferred to the full naturalistic stimulus set (9 continua × 128 morph steps; 1,152 syllables total), sampled linearly and equally spaced in latent space. Once performance was reliably above chance on every full continuum for several consecutive days, we introduced the cue syllables. When present, a cue syllable preceded the target by a 100-ms inter-syllable interval (Fig. 1A). The cue structure was designed to require integration of cue and target evidence rather than reliance on the cue alone: a cue preceded the target on 84% of trials (absent on 16%), of which 4% were uninformative (*p* = 0.5) and 80% were one of four informative cues: a “strong” (*p* = 0.875) and a “weak” (*p* = 0.75) cue per category, each on 20% of trials. Overall, 65% of trials were validly cued, 15% invalidly cued, 4% uninformatively cued, and 16% uncued. Because the cue was probabilistic, sometimes uninformative, and often absent, above-chance performance could not be achieved from the cue alone (Fig. 1A,B). For the present analyses we excluded uninformative (*p* = 0.5) cues and, within each category, pooled the strong and weak cues into a single cue, yielding three conditions per continuum: no cue, validly cued, and invalidly cued (pooled cue valid on ∼81% of cued trials). Subjects advanced to recording only after performing above 70% correct on the full cued task for 10k trials.

### Surgery and chronic electrophysiology

Surgical and chronic recording procedures were performed as described in [23]. We implanted subjects with a 32- or 64-channel NeuroNexus silicon probe, fitted on a custom 3D-printed micro-drive, targeting the auditory pallium. We acquired continuous extracellular waveforms across all probe channels at 30 kHz through a headstage (Intan Technologies) via OpenEphys acquisition hardware and software (Open-ephys.org). We recorded waveforms continuously over multiple days and operant behavioral blocks, synchronized with coincident trialized operant behavior and stimulus playback at sample-aligned precision. At the end of recording each day, passive playback blocks presented operant stimuli with the houselights off, the operant response ports inoperative, and the food hopper inaccessible, with a random 1.1 to 1.5 s inter-stimulus-interval.

### Spike sorting and single unit tracking

We extracted putative spiking activity from continuous extracellular waveforms in 12-h blocks using KiloSort (v2 and v2.5) [69] and SpikeInterface [70], after bandpass filtering (300–6,000 Hz) and common-median referencing [23]. A sorted cluster was screened with standard quality metrics (presence ratio, ISI-violation rate, SNR, and amplitude cutoff) and was retained as a single unit only if its mean template was a physiological, negative-going extracellular waveform and was spatially localized on the probe. We localized each putative unit to its peak (maximum amplitude) recording channel, whose anatomical position was derived from the probe geometry referenced against the stereotactic implant coordinates and an atlas of the starling brain [71]. Because the implanted arrays typically spanned two or more adjacent auditory regions, sorted individual units cannot be assigned to a single region with absolute certainty. Thus, our primary analyses pool units across the auditory forebrain. For the region-resolved comparison (Fig. S6) we assigned a neural population to a region only when at least 70% of its sorted units localized to that region by stereotaxic coordinates [23]. We present the region-resolved results as a best estimate of regional differences, but note that they are necessarily lower-powered because fewer recording days (neural populations) and subjects met the localization criterion.

### Neurophysiological Datasets

### Statistical analyses

Unless otherwise noted, data are reported as mean ± SEM. Throughout, “subject” denotes the animal and “neural population” the ensemble of units simultaneously recorded on a single recording day; our observations are hierarchically nested as trials within neural populations within subjects, with each subject contributing many recording days (range 1–59). We therefore evaluated every comparison with a linear mixed-effects model (LME) (statsmodels) carrying a by-subject random intercept and, where multiple observations per neural population entered the model, an additional by-population random intercept. This makes each fixed-effect estimate generalize across subjects rather than across non-independent neurons, recording days, or trials. The response variable and fixed-effect structure were chosen per analysis, and each reported effect is the corresponding fixed-effect estimate with its 95% confidence interval. Because the by-population and by-subject random structure can only be estimated where a penetration site contributes more than one analyzable population, we required each penetration site entering a model to have at least two analyzable recording days. As a subject-level cross-check we also computed one summary value per subject and tested it against the null with a paired *t* and Wilcoxon signed-rank test; where reported, the rank-biserial correlation *r* is the effect size for these descriptive cross-checks. Bootstrap 95% CIs were computed from *B* = 5000 resamples unless stated otherwise. Analyses were performed in Python 3.8.0 using NumPy (v1.24.3), SciPy (v1.10.1), statsmodels (v0.14.1), and scikit-learn (v1.3.2).

### Psychometric curve fitting

For each subject and cue condition (cue-left, cue-right, no-cue), behavioral responses were fit with a four-parameter logistic function of the form

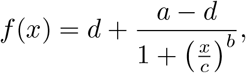

where *a* and *d* are the minimum and maximum asymptotes, *c* is the inflection point (the stimulus value at which *P* (right) = 0.5, i.e., the psychometric threshold *T*_50_), and *b* is Hill’s slope. Fits were performed on up to the most recent 100,000 trials per subject using `scipy.optimize.curve fit`. Cue conditions for a higher probability category-A cue versus a weaker category-A cue probability were pooled into a single category-A cue condition and the same for the two category-B cue conditions. Per-subject mean curves were computed by averaging fitted *f* (*x*) vectors across neural populations within each cue condition, after aligning orientation so that *P* (right) increases across the continuum. The psychometric midpoint *T*_50_ was extracted directly from the fitted inflection parameter *c*. Cue-dependent shifts in *T*_50_ were tested using two-sided paired Wilcoxon signed-rank tests across subjects (*n* = 10), with differences expressed as 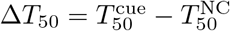.

### Spike vectors

Single-unit spike trains were converted to spike vectors following [23]. Spikes were binned into 20-ms time bins and smoothed with a Gaussian kernel (*σ* = 25 ms).

### Trial strata

The task used nine morph continua, each spanning 128 interpolation steps between category-A and category-B endpoints. Because individual morph steps yield few trials per condition, we grouped the 128 steps within each continuum into 4 bins of equal width, producing stimuli of similar perceptual difficulty. A *stratum* is defined as the intersection of one neural population, one morph continuum, and one difficulty bin, yielding a set of trials with comparable stimulus strength and shared recording context. Analyses requiring a minimum number of matched trials per condition (e.g., decoder fitting, cosine similarity estimation) use the stratum as the unit of observation; minimum trial thresholds are stated for each analysis.

### Within-category cosine similarity

For each neural population we compared responses to acoustically distinct, same-category target syllables (different signals on the same side of a continuum’s boundary, matched within continuum; see Trial strata), separately for valid and invalid cues. Single-neuron similarity was the mean pairwise cosine similarity between single-trial spike vectors of the two same-category stimuli, averaged across units. Population similarity was the cosine similarity between the PCA trajectories of the two stimuli. We used trial-count-matched subsamples (equal *n* per stimulus, averaged over *B* = 8 resamples) to remove the trial-number bias of averaged estimates. The single-neuron and population analyses were each fit with the same model,

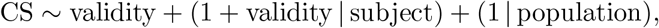

A one-sided per-subject paired Wilcoxon signed-rank test across populations was a cross-check at each scale.

### Dimensionality reduction with PCA

We constructed firing-rate estimates by binning spikes into 20-ms bins and smoothing with a Gaussian kernel (*σ* = 25 ms) [23]. For each neural population, we fit Principal Component Analysis on the trial-by-time-bin smoothed population spike train matrix with neurons as features [28], pooling all task conditions to obtain a shared subspace. To choose the number of retained components, we computed two complementary criteria per recording: the participation ratio, 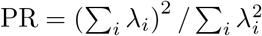 [35, 36], and the smallest *k* for which the cumulative variance explained reached 90%. Both criteria converged on *K* = 8 as the rounded median across recordings, which we used for all subsequent population-level analyses.

### Stimulus Identity Decoder

Stimulus identity was decoded from the single-trial population state with a cross-validated linear classifier (shrinkage linear discriminant analysis, stratified 5-fold cross-validation, balanced accuracy). Two target factors were decoded separately: the continuum identity (nine continua) and the bin along the continuum (four bins). Chance was estimated empirically by label permutation, and the per-population gain over chance (Δ balanced accuracy) was tested with a linear mixed model with a by-subject random intercept, matching the inference used for the cosine-similarity analyses.

### Degeneracy-enabled remapping latent-dynamics model

All model panels (Fig. 3A–J) were run with (`numpy` default generator, seed = 22): *N* = 100 model neurons, *K* = 3 latent dimensions, *T* = 60 bins of 20 ms, target-stimulus window [*t*_0_, *t*_1_) = [100ms, 1100ms), and *N*_trials_ = 600 trials per condition.

#### Latent state and readout

The latent state *x*_*t*_ ∈ ℝ^*K*^ evolves as in Eq. (1), with the fixed asymmetric transition matrix

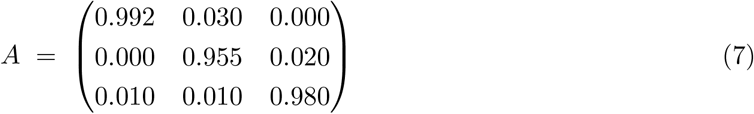

(eigenvalues ≈ 1.000, 0.973, 0.953). The drive *u*_*t*_ is non-zero only during the target-stimulus window, and the dynamics noise *σ*_*t*_ switches from *σ*_pre_ = 0.010 to 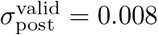 or 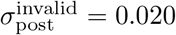 at target onset. Firing rates follow the linear readout of Eqs. (2) – (4) (no rectification); the baseline *r*_0_ = 13 and gain *g* = 1.2 are fixed and serve only to keep rates non-negative, so neither affects the mean-centered geometry. The rotation time constant was *τ*_rot_ = 200 ms. A Gaussian base readout *W*_base_ ∈ ℝ^*N ×K*^ (scaled by 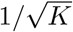) gave the pre-target readout *W*_pre-target_ and post-target template 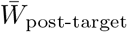 by rotating its column span by *θ*_prep_ = 4^*°*^ and *θ*_target_ = 6^*°*^ in random orthogonal planes.

#### Task-potent axis and null/potent projectors

The task-potent direction of Eq. (5) is the category-readout axis

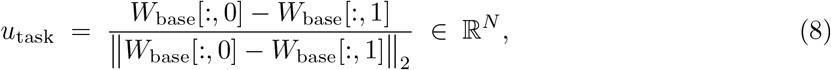

with projectors 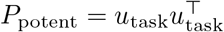 and *P*_null_ = *I*_*N*_ −*P*_potent_. We use its latent preimage *u*_pot-latent_ ∝ 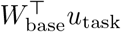 and a fixed residual latent direction *v*_res_ orthogonal to it.

#### Identity remap with null/potent split

The trial-specific post-target readout is 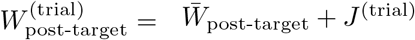, where the remap splits into independent null and potent contributions,

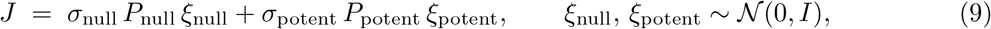

following [7, 11, 13]. Separate *σ* values set the relative magnitude of behaviorally-invisible and behaviorally-consequential variability per condition. Each trial’s 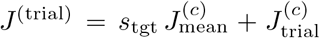 condition mean plus per-trial jitter (Table 2). On trials with a valid cue, a fixed per-neuron sign vector *s*_*n*_ ∈ {+1, −1}^*N*^ is applied, decorrelating a neuron’s tuning across targets; on trials with an invalid cue *s*_tgt_ = ±1 matches the target sign, breaking that antipodal symmetry. The result is null-dominated remapping on trials with a valid cue and potent-dominated remapping on trials with an invalid cue.

**Table 2:** Identity-remap magnitudes for the null and potent components of *J* (Eq. (9); seed = 22).

|  | condition mean |  | per-trial jitter |  |
| --- | --- | --- | --- | --- |
| | $\sigma_{\text{null}}$ | $\sigma_{\text{potent}}$ | $\sigma_{\text{null}}$ | $\sigma_{\text{potent}}$ |
| Valid | 0.20 | 0.005 | 0.160 | 0.001 |
| Invalid | 0.02 | 0.30 | 0.005 | 2.0 |

#### Per-trial trajectory rotation

We measured the angle between a pre-target anchor (mean over [0, *t*_0_)) and the state at each bin,

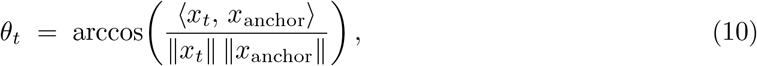

averaging over [*t*_0_, *t*_1_) for stim-window summaries.

#### Per-neuron weight displacement

Per neuron we took 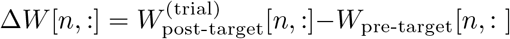 and split it as in Eq. (6). Fig. 3D plots ∥Δ*W*_null_[*n*, :]∥_2_ versus ∥Δ*W*_potent_[*n*, :]∥_2_; Fig. 3E plots the null fraction

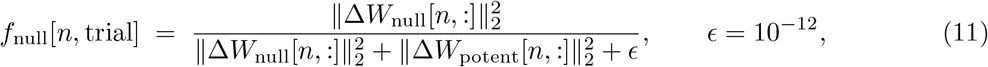

which is 1 for a purely null and 0 for a purely potent displacement.

#### Cosine similarity

For single neurons, we took the mean off-diagonal pairwise cosine of each trial’s target-window firing trace across trials within each validity and target pairing, averaging the two targets to give one value per neuron. For populations, we projected each trial’s target-window activity into the top-*K* (*K* = 3) principal-component subspace of the pooled population activity and took the mean off-diagonal pairwise cosine within each validity and target) pairing.

### Empirical subspace geometry analyses

#### Task-potent decision axis

Several analyses share a single population-level task-potent axis, computed two ways depending on whether the trials being scored must be held out from the fit. In both cases, for each neural population an L2-regularized logistic regression (`scikit-learn`, `liblinear`, *C* = 1.0, class-balanced) was fit on stimulus-window mean activity to discriminate category-A from category-B (left-vs. right-port) targets; the unit-normalized weight vector **w** ∈ ℝ^*N*^ defines the task-potent axis *U* = **w**, with potent projector *P*_potent_ = *UU*^*⊤*^ and its orthogonal complement as the null subspace. For the null-fraction and rotation analyses (Fig. 3K–N), where the axis only summarizes a within-condition readout direction, the decoder was fit on correct active trials pooled across cue validity (minimum 10 trials per side)

#### Null-subspace reorganization of target-evoked activity

For every trial we took the stimulus-window mean population vector relative to the pre-target mean and split this displacement into its potent component (along *U*, via *P*_potent_) and its null remainder. Each trial was summarized by its null fraction, the share of the displacement’s total squared magnitude in the null subspace,

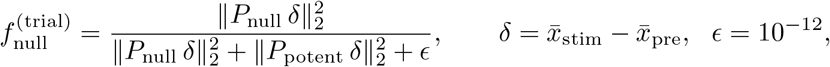

computed over the whole population. Trial null fractions were averaged within each population × stimulus stratum (minimum 5 trials per validity) and then to one value per neural population and validity. The same decomposition was applied to passive-playback trials (Fig. 3M). LME (Fig. 3K, active): *f*_null_ ∼ validity + (1 | subject) + (1 | population), *β* = valid − invalid, one-sided alternative valid *>* invalid; corroborated by a per-subject paired Wilcoxon. The passive analysis (Fig. 3M) used the identical model on passive trials.

#### Pre-to-stimulus subspace rotation

For each population we defined a pre-target subspace as the top *k* = 3 principal components of trial-averaged pre-target activity, and a stimulus subspace at each bin as the top *k* = 3 PCs of trial-averaged activity in a symmetric ±40 ms window centered on that bin (matched to the length of the pre-target stimulus window, so the two bases are estimated from equal amounts of data). Reorientation was quantified as the mean canonical principal angle between the pre-target and stimulus bases,

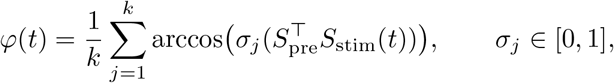

yielding a rotation time course per (neural population, target, validity) stratum (minimum 5 trials); target stimulus window means were averaged to one value per neural population and validity. Robustness across *k* = 2–6 is shown in Fig. S4, and the maximum principal angle gives the same ordering. The same analysis was applied to passive trials (Fig. 3N). LME (Fig. 3L, active): *φ* ∼ validity + (1 | subject) + (1 | population), *β* = invalid − valid, one-sided alternative invalid *>* valid; per-subject paired Wilcoxon as a cross-check. The passive analysis (Fig. 3N) used the identical model on passive trials.

#### Noise correlation

Within each population × stimulus stratum we formed the trial × neuron matrix of target-stimulus window time-averaged activity, converted its neuron-by-neuron sample covariance to a correlation matrix, and took the mean of its off-diagonal pairwise noise correlations. Stratum values were averaged to one value per neural population and validity. 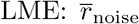, *β* = valid − invalid, one-sided alternative valid *>* invalid.

#### Cross-validated PCA and effective dimensionality

For each neural population and validity we pooled active correct trials and computed a cross-validated eigenspectrum [4]: over *n*_splits_ = 50 random half-splits (seed 0) we estimated principal components on one half, centred the held-out half with the *training* mean, projected it onto the training PCs, and took the held-out variance per component; these were normalized to sum to one and averaged across splits, up to *K*_max_ = 20 modes. The participation ratio was 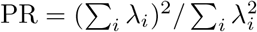 [35, 36] computed on the cross-validated spectrum. LME (Fig. 3P, per mode): for each mode *k*, 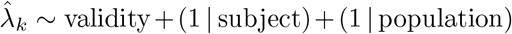, *β* = valid − invalid, with Benjamini–Hochberg FDR correction across the 20 modes; markers in Fig. 3P denote FDR-corrected significance. LME (Fig. 3Q, effective dimensionality): PR ∼ validity+ (1 | subject)+(1 | population), *β* = valid−invalid; a per-subject paired Wilcoxon on the participation ratio served as a cross-check.

#### Held-out (no-cue-trained) task-potent axis

For the margin analyses (Fig. 4A,B) the same axis was instead fit on no-cue correct trials only (minimum 10 trials per side) and then applied to the held-out cued (valid and invalid) trials. Training on no-cue trials keeps the axis independent of the cued trials it scores, so the cue-validity (Fig. 4A) and accuracy (Fig. 4B) contrasts are not circular. Each cued trial’s signed decision-function value (margin) was flipped so that positive values point toward the trial’s true target category; this held-out axis is the empirical analog of the model’s *u*_task_ and is shared by the directional-shift and wrong-side analyses below.

#### Directional task-potent shift

On every cued trial we took the target-signed margin along this axis. Within each population, margins were averaged within target strata (minimum 3 trials with a valid cue and 3 trials with an invalid cue per stratum) and then across strata, removing stimulus-composition and valid/invalid imbalance. LME: margin ∼ validity + (1 | subject) + (1 | population), with *β* = valid − invalid; a one-sided paired Wilcoxon signed-rank test across populations was a cross-check.

#### Wrong-side decoder margins

Using the same axis, every active trial’s signed margin was scored as correct- or wrong-side (negative = wrong side). Per population we computed the wrong-side fraction for correct and incorrect trials, balanced within stimulus strata (minimum 3 trials per accuracy class per stratum). LME: wrong-side ∼ accuracy + (1 | subject) + (1 | population), *β* = incorrect − correct; corroborated by a one-sided paired Wilcoxon across populations.

#### Reaction time across cue conditions

Reaction time (RT) was the interval between target stimulus offset and the pecking response (trials rewarded only if the subject waited until target offset). The primary test treated the subject as the unit of analysis. Per-subject median RTs for valid, invalid, and no-cue trials were compared with a Friedman test and pairwise Wilcoxon signed-rank tests. This was corroborated by a subject-aware trial-level linear mixed-effects model, RT ∼ validity + (1 | subject).

#### Pre-target centroid geometry

Pre-target activity vectors (correct trials) were *z*-scored within trial. Per population (minimum 10 trials per group) we computed pre-target centroids for category-A cue, category-B cue (trials with a valid cue), and no-cue trials, and the displacement ratio

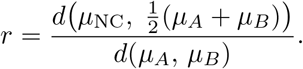

The geometry was tested with an intercept-only LME on the raw displacement difference, *d*(*µ*_NC_, mid) − *d*(*µ*_*A*_, *µ*_*B*_) ∼ 1 +(1 | subject), against zero; a one-sided signed-rank test on (*r* − 1) *>* 0 and a 10,000-replicate bootstrap CI on the median were cross-checks.

#### Trajectory distance from pre-target subspace

A *K* = 3 PCA subspace *Q*_pre_ was estimated from trial-averaged pre-stimulus activity. At each stimulus-epoch bin the distance of the running mean vector from this subspace,

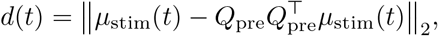

was normalized by the median norm of pre-stimulus trial residuals. Cued and no-cue curves were paired within population (stimulus-window mean per curve). Active trials were tested with Δ_cued*−*NC_ ∼ 1 + (1 | subject); the active-versus-passive comparison used populations recorded in both modalities, distance ∼ cue×modality +(1 | subject) +(1 | population), with the cue × modality interaction as the test.

#### Trial-by-trial pre-target geometry and reaction time

For each active correct trial (RT 0.05– 5 s) we computed the peak distance of the running stimulus-window mean from the trial’s cue-group pre-target subspace (top *k* = 3 pre-target PCs, maximized across expanding windows). The metric was *z*-scored within neural population and related to log RT with log RT ∼ distance +(1 | subject) + (1 | population), where *β* is the change in log RT per +1 SD; pooled and per-population (≥ 30 trials) Spearman correlations were cross-checks. Pre-target variance and distance from the population centroid (Fig. S5) were analyzed identically.

## SUPPLEMENTARY

**Table S1:** Per-subject psychometric parameters. Psychometric midpoint *T*_50_ (continuum steps) and Hill slope from four-parameter logistic fits, per subject and cue condition (*n* = 10 subjects). *n* trials is the total across the three cue conditions.

| Subject | $n$ trials | $T_{50}$ (steps) | | | Slope | | |
| --- | --- | --- | --- | --- | --- | --- | --- |
|  |  | No cue | cue-A | cue-B | No cue | cue-A | cue-B |
| B1170 | 95,852 | 62.2 | 58.4 | 66.4 | 8.6 | 9.2 | 9.2 |
| B1248 | 88,144 | 60.7 | 65.9 | 58.1 | 7.0 | 8.5 | 7.0 |
| B1597 | 32,006 | 61.7 | 59.9 | 68.7 | 16.6 | 17.4 | 10.8 |
| B1595 | 90,254 | 63.4 | 69.9 | 63.4 | 12.3 | 9.1 | 8.8 |
| B1188 | 73,345 | 68.8 | 60.2 | 65.4 | 11.9 | 6.0 | 7.3 |
| B1244 | 96,086 | 61.7 | 58.5 | 67.1 | 9.1 | 10.7 | 9.2 |
| B1276 | 95,920 | 65.7 | 67.7 | 63.3 | 11.6 | 13.8 | 13.8 |
| B1426 | 95,984 | 62.2 | 57.4 | 67.8 | 11.4 | 9.5 | 10.0 |
| B1432 | 96,002 | 63.1 | 61.1 | 65.1 | 28.8 | 25.8 | 26.6 |
| B1593 | 95,993 | 60.9 | 66.8 | 57.9 | 8.4 | 10.5 | 9.7 |

**Figure S1:**
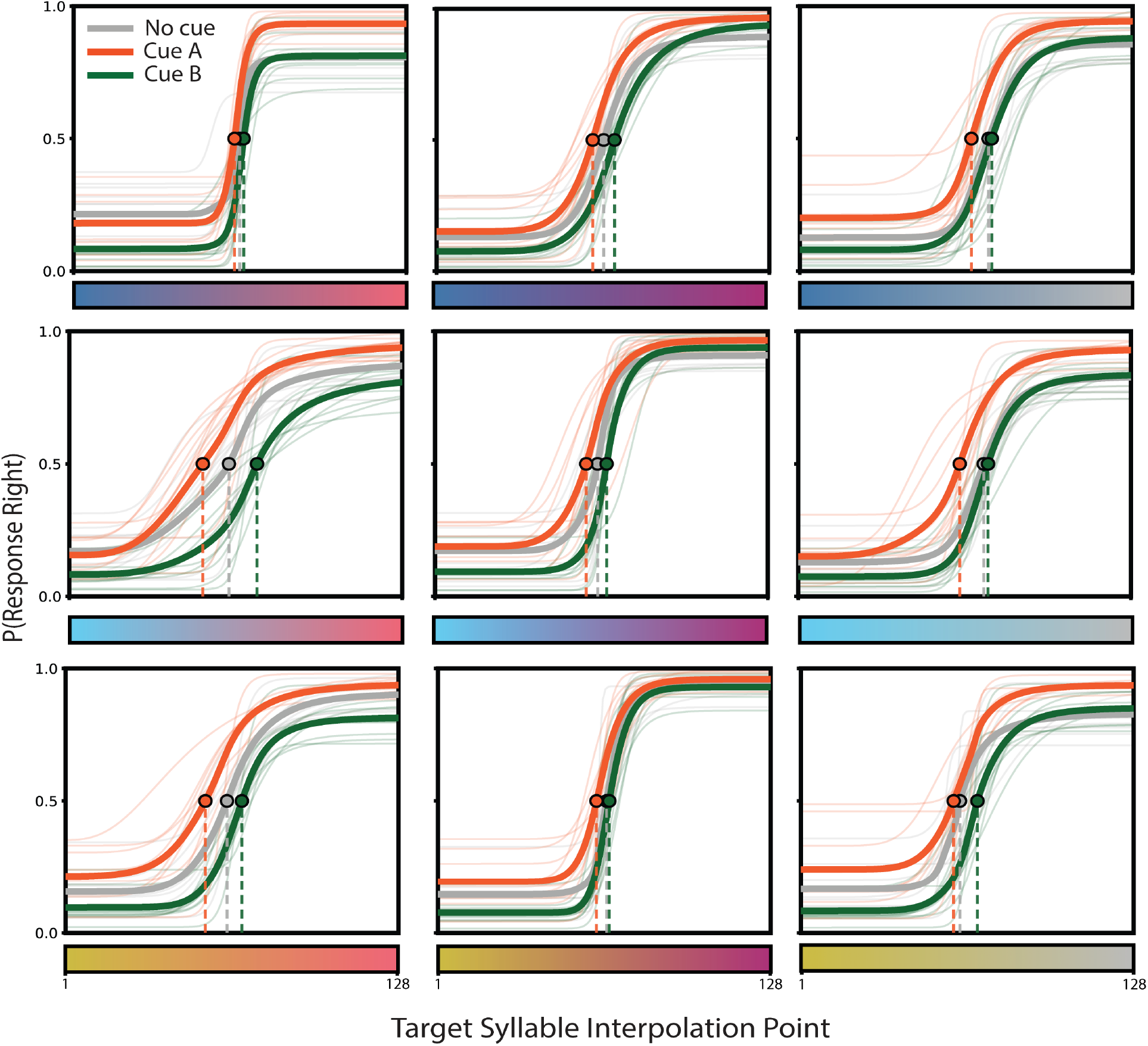
Cue-conditioned psychometric functions are consistent across all nine syllable continua. Each panel is one stimulus continuum, identified by its endpoint colors: the left endpoint sets the row (blue, top; cyan, middle; gold, bottom) and the right endpoint sets the column (red, left; purple, middle; grey, right). Curves are four-parameter logistic fits of *P* (peck right) versus position along the morph continuum (1 = left endpoint, 128 = right endpoint), fit separately for No cue, cue-A, and cue-B trials. Thin lines are individual subjects; thick lines are the across-subject mean; filled circles mark the 50% threshold (*T*_50_) of the mean curve with a dropline to the abscissa. Per-continuum values in Table S2). Only (subject, continuum, cue) cells with *≥*50 trials were fit.

**Table S2:** Cue-induced shifts in categorization threshold across syllable continua. Median change in the psychometric midpoint, Δ*T*_50_ (continuum steps), relative to no-cue trials, for category-A cues and category-B cues. Positive values indicate a shift toward category A and negative values a shift toward category B; that is, each cue shifts the threshold toward its predicted category, matching the convention in Fig. 1D. Continua are named by their endpoint colors (left *→* right), matching the panels in Fig. S1. *p*-values are two-sided paired Wilcoxon signed-rank tests across subjects (per-continuum *n* = subjects; tests uncorrected). The pooled row tests all subject × continuum observations (*n* = 85 per contrast; Wilcoxon *W* = 302 for the cue-A contrast, *W* = 245 for the cue-B contrast).

| Continuum | *n* | cue-A – no cue |  | cue-B – no cue |  |
| --- | --- | --- | --- | --- | --- |
| | | $\Delta T_{50}$ | *p* | $\Delta T_{50}$ | *p* |
| Blue→Red | 10 | +1.95 | 0.084 | −1.53 | 0.0020 |
| Blue→Purple | 10 | +5.06 | 0.0020 | −5.23 | 0.0020 |
| Blue→Grey | 9 | +6.97 | 0.0039 | −2.32 | 0.25 |
| Cyan→Red | 10 | +9.05 | 0.020 | −17.7 | 0.0020 |
| Cyan→Purple | 10 | +4.16 | 0.084 | −4.71 | 0.0020 |
| Cyan→Grey | 9 | +8.60 | 0.0039 | −2.91 | 0.16 |
| Gold→Red | 9 | +8.96 | 0.0078 | −5.53 | 0.0039 |
| Gold→Purple | 9 | +4.63 | 0.027 | −0.95 | 0.30 |
| Gold→Grey | 9 | +5.13 | 0.098 | −4.76 | 0.0039 |
| Pooled | 85 | +4.97 | $2 \times 10^{-11}$ | −3.73 | $4 \times 10^{-12}$ |

**Figure S2:**
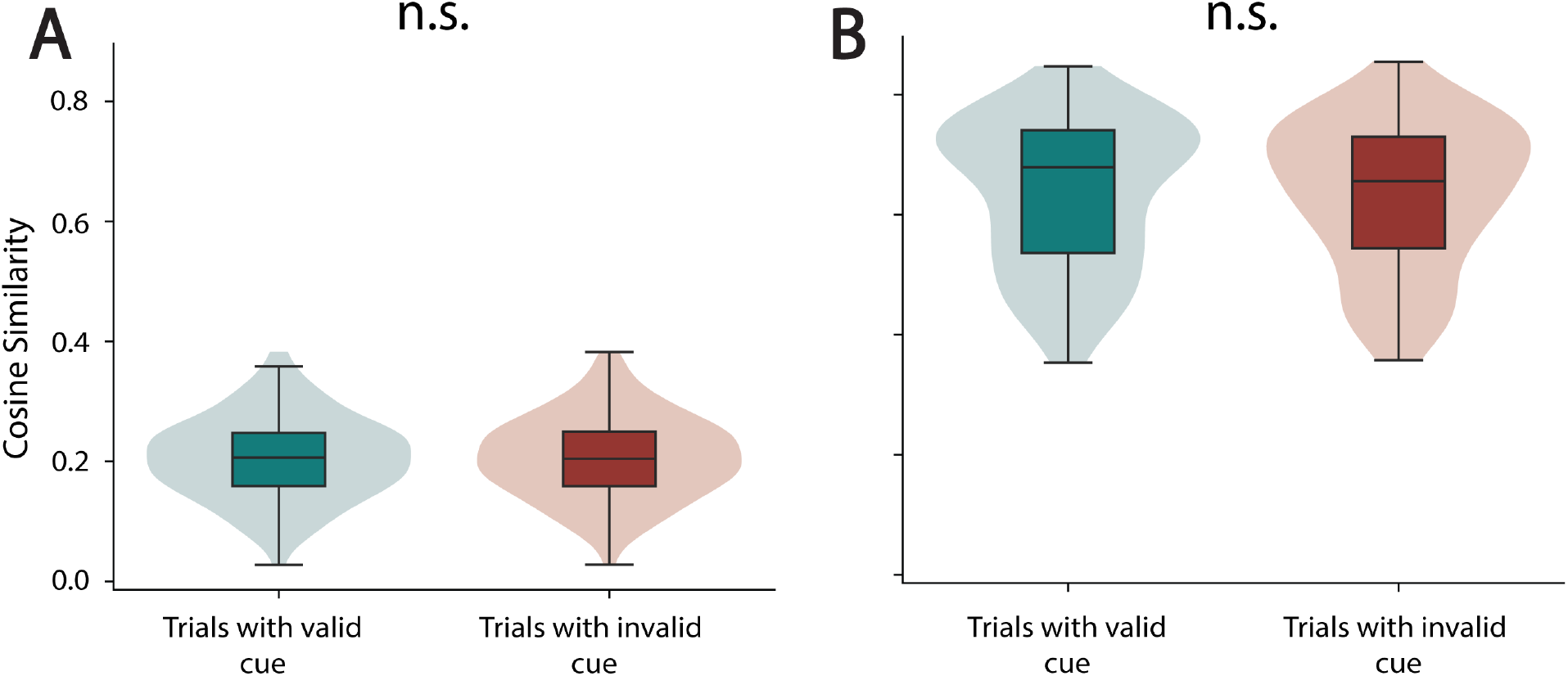
Within-category cosine similarity is unaffected by cue validity during passive playback. Boxes show the population-level distribution (shaded violin) with the median and interquartile range. **(A)** Single-neuron within-category cosine similarity (*n* = 108 neural populations from 5 subjects). **(B)** Population within-category cosine similarity (*n* = 109 neural populations from 6 subjects).

**Figure S3:**
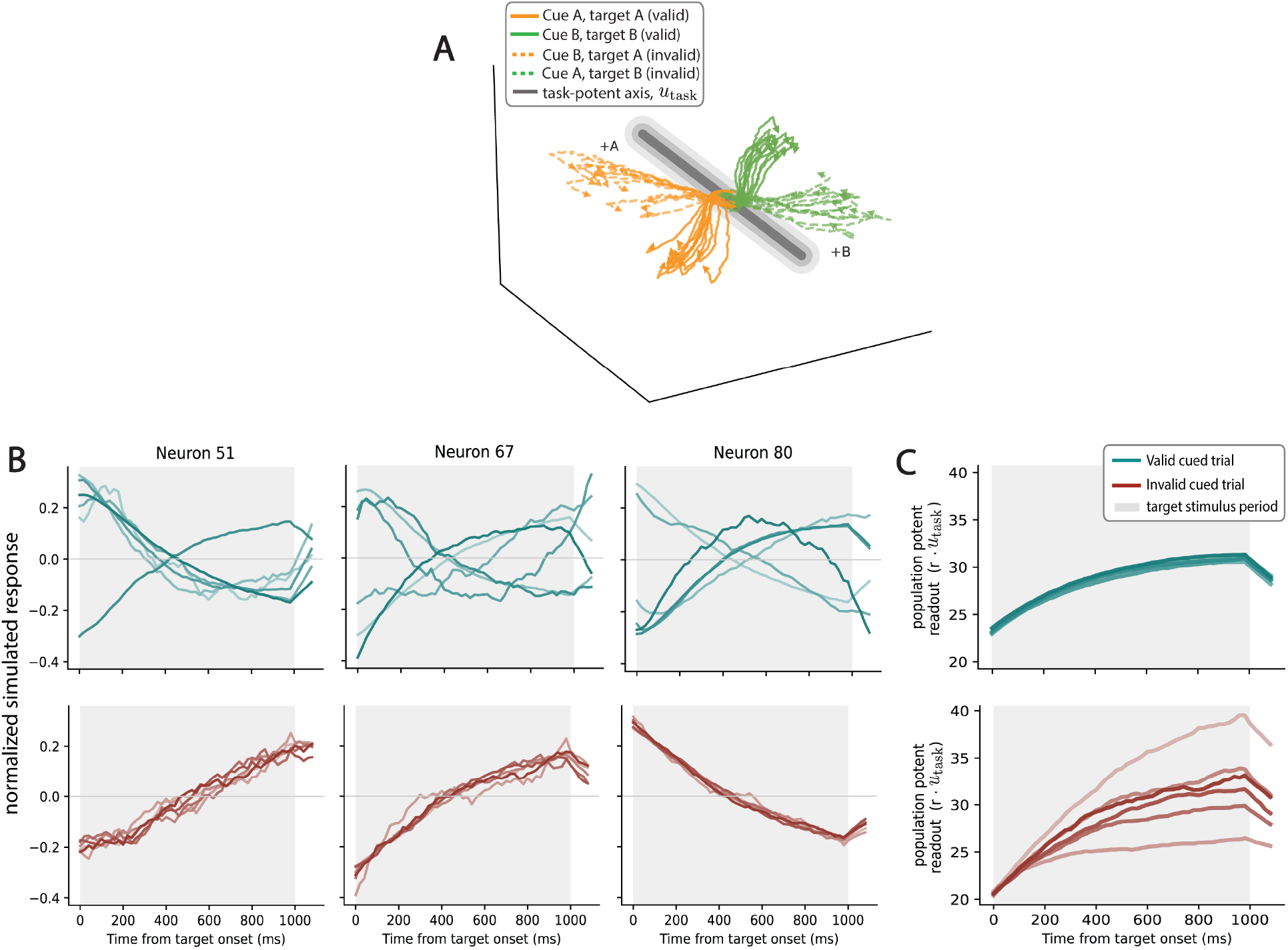
Degeneracy-enabled remapping: divergent single-neuron responses project onto a shared task-potent population readout. **(A)** Three-dimensional PCA of simulated population trajectories under valid and invalid cues, colored by target category (solid lines, valid; dashed lines, invalid). The task-potent axis *u*_task_ is the direction along which category-A and category-B mean responses differ. We project this into the PCA space and drawn as the gray line, with its category-A (+A) and category-B (+B) ends labeled. The coding axis is a single direction within the full population space; valid trajectories settle tightly toward its ends while invalid trajectories scatter along it (*n* = 12 trials per condition). **(B)** Normalized single-neuron responses for three example neurons on six valid-cue trials (top row) and six invalid-cue trials (bottom row) trials to the same target. Each trace is one trial’s stimulus-window response, unit-normalized. The shaded region marks the target stimulus period. **(C)** Population readout, the projection of population activity onto the task-potent axis (*r u*_task_), for the same six valid (top) and six invalid (bottom) trials. Despite the heterogeneous single-neuron responses in (B), the valid-cued trials readout collapses onto a nearly identical trajectory across trials, whereas the invalid-cued trials readout is dispersed across trials.

**Figure S4:**
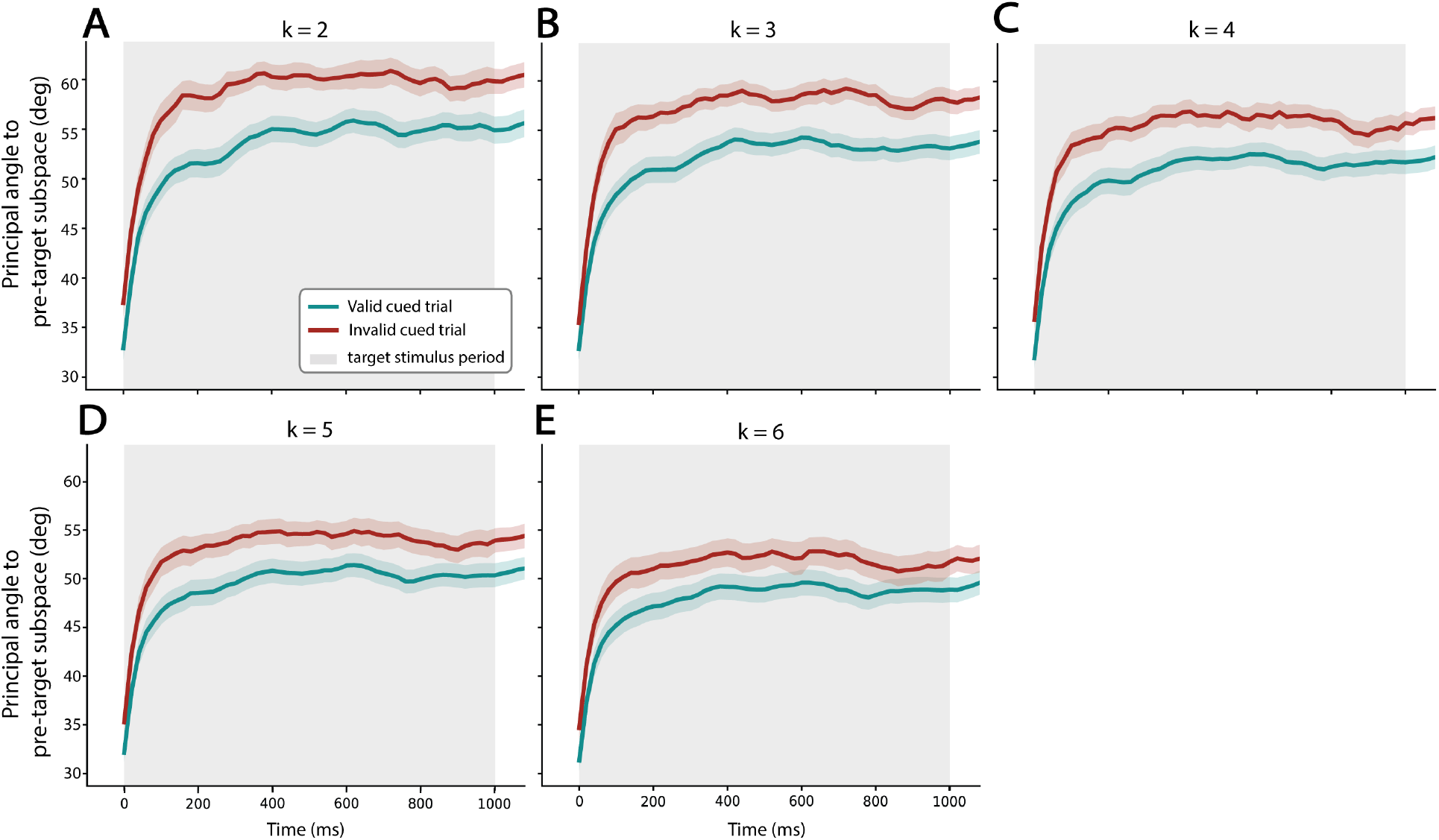
Pre-to-stimulus subspace rotation is robust across choices of subspace dimensionality *k*_sub_. Each panel shows the rotation timecourse (mean *±* SEM across neural populations) for a different *k*_sub_; rotation is the mean principal angle (degrees) between the pre-target subspace and the current subspace (top *k*_sub_ PCA components). **(A)** *k*_sub_ = 2: *β* = +6.00^*°*^, 95% CI [+4.16^*°*^, +7.84^*°*^], *p* = 1.7 *×* 10^*−*10^. **(B)** *k*_sub_ = 3: *β* = +5.67^*°*^, 95% CI [+4.11^*°*^, +7.23^*°*^], *p* = 1.1 *×* 10^*−*12^. **(C)** *k*_sub_ = 4: *β* = +5.23^*°*^, 95% CI [+3.69^*°*^, +6.77^*°*^], *p* = 3.0 *×* 10^*−*11^. **(D)** *k*_sub_ = 5: *β* = +4.75^*°*^, 95% CI [+3.21^*°*^, +6.30^*°*^], *p* = 1.7 *×* 10^*−*9^. **(E)** *k*_sub_ = 6: *β* = +4.26^*°*^, 95% CI [+2.82^*°*^, +5.71^*°*^], *p* = 7.4 *×* 10^*−*9^. The invalid *>* valid effect was preserved across all *k*_sub_ tested and largest at the lowest *k*_sub_ and attenuating as *k*_sub_ grew consistent with the headline rotation analysis at *k*_sub_ = 3 (Fig. 3L).

**Figure S5:**
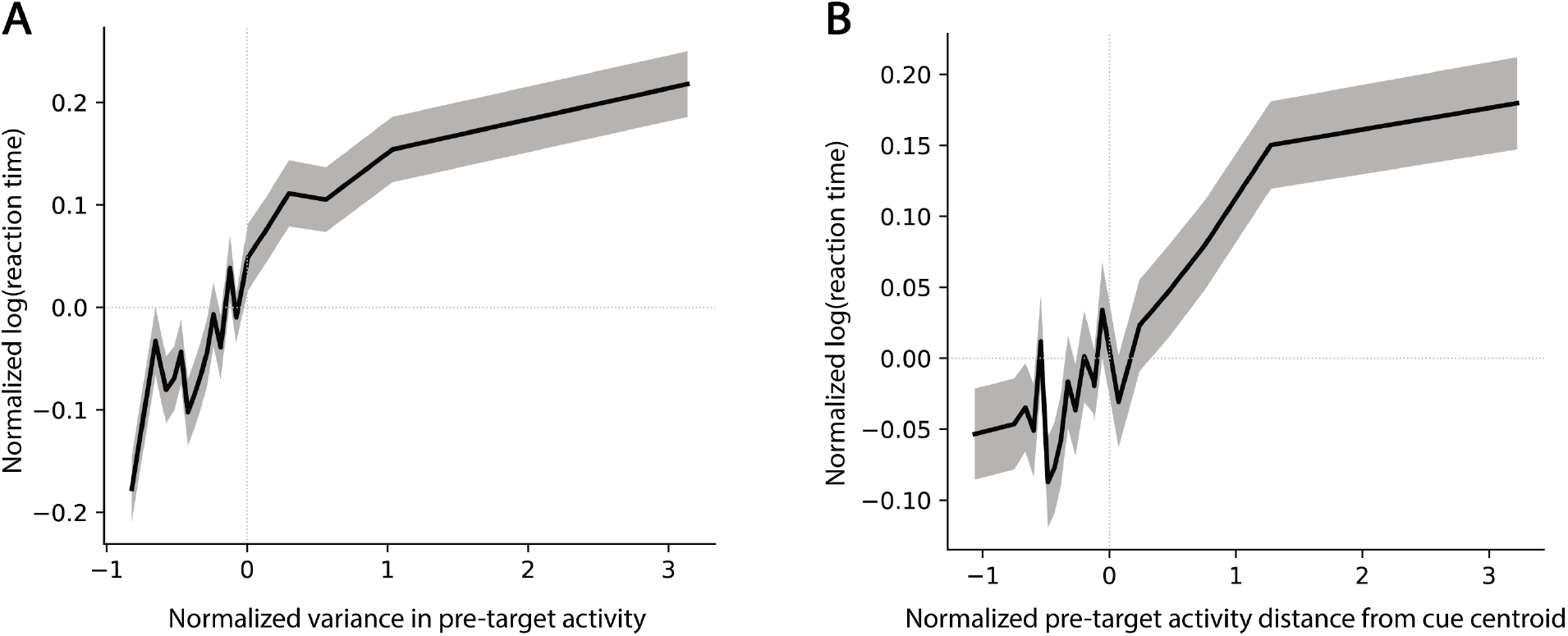
Pre-target stability predicts reaction time under alternative operationalizations. Both panels show within-population z-scored pre-target stability versus log reaction time, pooled across all active correct trials (*n* = 89,232 trials, *n* = 132 neural populations, 7 subjects). Black lines show mean log(reaction time) across 20 equal-frequency bins; shading represents *±*SEM. **(A)** Pre-target neural variance. Trials with greater pre-target variance had slower reaction times (pooled Spearman *ρ* = +0.063, *p* = 3.9 *×* 10^*−*78^; median per-population *ρ* = +0.053; *p* = 1.7 *×* 10^*−*11^). **(B)** Distance from pre-target cue centroid. Trials farther from the population-typical pre-target state had slower reaction times (pooled Spearman *ρ* = +0.049, *p* = 1.0 *×* 10^*−*48^; median per-population *ρ* = +0.029; *p* = 5.4 *×* 10^*−*9^). Together, these results confirm that the relationship between pre-target population stability and reaction time is robust to the choice of stability measure.

**Figure S6:**
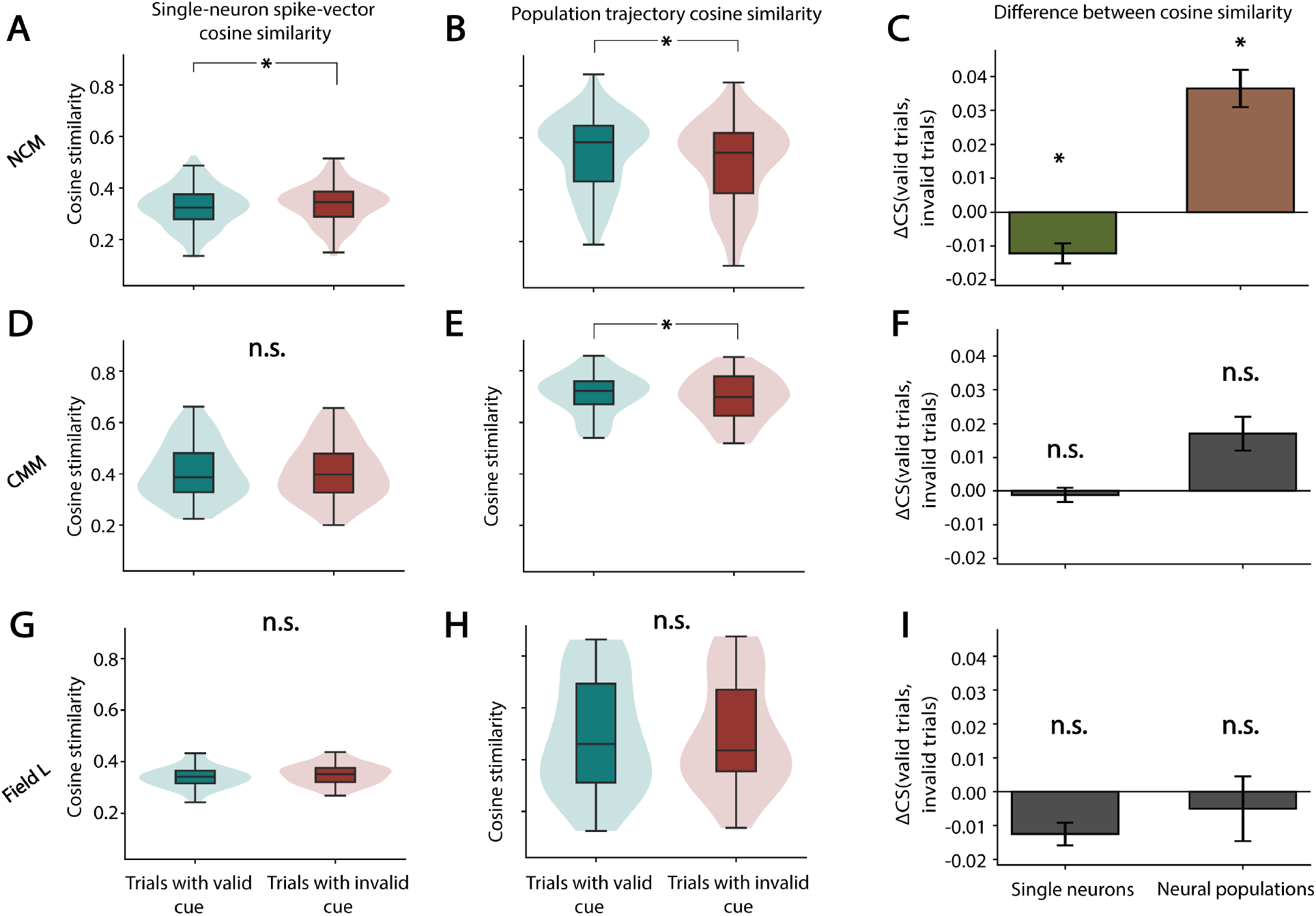
Cue validity effects on single-neuron and population-level stimulus encoding across the auditory hierarchy. All effects are linear mixed-effects estimates of the valid invalid difference with a by-subject random intercept, evaluated at the level of the neural population. In the violin panels (A, B, D, E, G, H), each violin shows the distribution across neural populations (box, interquartile range; central line, median); in the bar panels (C, F, I), bars show the mean ΔCS (valid *−* invalid) *±*SEM. * denotes a significant valid–invalid difference; n.s., not significant. **(A)** Single-neuron spike-vector cosine similarity in NCM (*n* = 123 neural populations from 3 subjects; *β* = *−*0.013, *p* = 2.9 × 10^*−*3^). **(B)** Population trajectory cosine similarity in NCM (*n* = 127 neural populations from 3 subjects; *β* = +0.045, *p* = 4.4 × 10^*−*3^). **(C)** Mean ΔCS (valid *−* invalid) in NCM for single neurons (*β* = *−*0.013, *p* = 2.9 × 10^*−*3^) and neural populations (*β* = +0.045, *p* = 4.4 × 10^*−*3^); both the single-neuron and population effects are present. **(D)** Single-neuron spike-vector cosine similarity in CMM (*n* = 44 neural populations from 3 subjects; *β* = +0.015, *p* = 0.31, n.s.). **(E)** Population trajectory cosine similarity in CMM (*n* = 41 neural populations from 3 subjects; *β* = +0.016, *p* = 0.57, n.s.). **(F)** Mean ΔCS (valid *−* invalid) in CMM for single neurons (*β* = +0.015, *p* = 0.31, n.s.) and neural populations (*β* = +0.016, *p* = 0.57, n.s.). **(G)** Single-neuron spike-vector cosine similarity in Field L (*β* not computable; *n* = 1 subject, not testable with a by-subject model and reported descriptively). **(H)** Population trajectory cosine similarity in Field L (*β* not computable; *n* = 2 subjects, not testable with a by-subject model and reported descriptively). **(I)** Mean ΔCS (valid *−* invalid) in Field L for single neurons and neural populations (*β* not computable); with only one to two subjects, neither scale is testable with a subject-level model, so values are descriptive only. Across regions, the population-level cue effect holds within NCM but not CMM, and single-neuron sharpening reaches significance within NCM as well as when pooled across the auditory forebrain (main Fig. 2); within CMM neither scale is individually significant, and Field L, with only one to two subjects, cannot be tested.

